# Inhibitory midbrain neurons mediate decision making

**DOI:** 10.1101/2020.02.25.965699

**Authors:** Jaclyn Essig, Joshua B. Hunt, Gidon Felsen

## Abstract

Decision making is critical for survival but its neural basis is unclear. Here we examine how functional neural circuitry in the output layers of the midbrain superior colliculus (SC) mediates spatial choice, an SC-dependent tractable form of decision making. We focus on the role of inhibitory SC neurons, using optogenetics to record and manipulate their activity in behaving mice. Based on data from SC slice experiments and on a canonical role of inhibitory neurons in cortical microcircuits, we hypothesized that inhibitory SC neurons locally inhibit premotor output neurons that represent contralateral targets. However, our experimental results refuted this hypothesis. An attractor model revealed that our results were instead consistent with inhibitory neurons providing long-range inhibition between the two SCs, and terminal activation experiments supported this architecture. Our study provides mechanistic evidence for competitive inhibition between populations representing discrete choices, a common motif in theoretical models of decision making.

## Introduction

Decision making is fundamental for adaptive behavior and examining its neural bases, along with developing corresponding theoretical frameworks, offers insight into circuit mechanisms for cognitive processes^1–3^. The potential utility of theoretical models is limited, however, by the extent to which we understand how the computations that give rise to decisions are implemented by circuits in the brain^4^. Recent technologies facilitating cell-type-specific examination of neural circuits in vivo have begun to close the sizeable gap between circuit computations and behavior^5^. Elucidating the functional circuitry underlying decision making is therefore a fruitful approach for uncovering the neural mechanisms of cognition.

An effective strategy to study decision making circuits is to examine neural activity in highly conserved brain areas during ethologically-relevant and computationally-tractable behaviors^6^. Spatial choice – selecting where to attend or to move – is ideal for this purpose: it is an adaptive form of decision making that is critical for survival, and several lines of evidence, across primates, cats and rodents, implicate the intermediate and deep layers of the midbrain superior colliculus (SC) in this function^7–9^. Targets in contralateral space are represented in the SC earlier than in other choice-related brain regions^10–15^, and manipulating SC activity during choice produces predictable choice biases^15–20^. These and other data showing that SC activity reflects target value and other decision-related variables^12, 21, 22^ demonstrate that the SC, traditionally appreciated for initiating orienting motor commands, is required for choosing where to move^7^. While the SC has recently served as an exquisite experimental system for probing neural circuit function in multiple behavioral contexts^23–26^, how SC circuits mediate spatial choice remains unclear.

The wealth of neural and behavioral data acquired from the SC during perceptual decision-making tasks has served as a foundation for developing theoretical models of decision making^27–32^. Inhibition is a critical feature of many of these models^30–32^. However, the functional role of inhibitory neurons – which comprise approximately 30% of SC neurons^33, 34^ – has been difficult to dissociate from the role of extrinsic inhibition^30, 35^, due to the inability of traditional methods to target GABAergic neurons for recording and manipulation. We therefore lack a clear understanding of how the intrinsic inhibition posited by these models is implemented, limiting their utility for elucidating neural circuit function. Anatomical and physiological data from rodent slice experiments suggest a role for GABAergic SC neurons in local inhibition of neighboring neurons^36, 37^, although evidence for long-range intercollicular inhibition has also been described in rodents and other species^34, 38, 39^. Ultimately, characterizing the functional connectome of the SC is insufficient for understanding the role of inhibitory SC neurons in decision making. To address this question, the activity of inhibitory SC neurons must be examined while the circuit is naturally engaged; i.e., during goal-directed behavior^5, 6^. Thus, we sought to determine the role of inhibitory SC neurons in mediating spatial choice by employing optogenetic strategies to manipulate and record their activity in behaving mice.

Given the data from rodent SC slice studies^37^ and the canonical role of GABAergic neurons in providing local inhibition in cortical circuits during perceptual decision-making^40^, we hypothesized that GABAergic SC neurons would inhibit nearby premotor output neurons during spatial choice. We predicted that, since premotor output neurons encode contralateral choice, activating inhibitory SC neurons during decision making would decrease the likelihood of selecting the contralateral choice, resulting in an ipsilateral choice bias. Surprisingly, we found the opposite effect—activating GABAergic neurons produced a contralateral choice bias. We followed up on these results with cell-type-specific recordings during behavior to reveal a direct relationship between GABAergic activity and contralateral choices. Finally, we developed a biologically-constrained attractor model to examine mechanisms by which GABAergic SC neurons could promote contralateral choices. Surprisingly, the model predicted a role for intercollicular inhibition in spatial choice for which we subsequently found experimental support. Overall, we have elucidated the function of inhibitory neurons in an important decision-making circuit and these results will help to inform our theoretical understanding of decision-making circuits in general.

## Results

### Activating GABAergic SC neurons promotes contralateral choices

We aimed to identify the functional role of inhibitory neurons in the output (intermediate and deep) layers of the superior colliculus (SC) in spatial decision making. We hypothesized that these neurons function to locally inhibit premotor output neurons (Fig. 1a). We first tested this hypothesis by selectively expressing channelrhodopsin (ChR2) unilaterally in GABAergic SC neurons of Gad2-Cre mice (Methods) and examining how photoactivating these neurons affected neural activity in neighboring SC neurons (Methods). Consistent with our hypothesis, we found that a subset of neurons exhibited a decrease in activity upon activation of adjacent GABAergic neurons (Fig. 1b; 32/301 neurons inhibited during stimulation/recording sessions; *p* < 0.05, one-tailed Wilcoxon signed-rank test; Methods), confirming the presence of local inhibition. To test our hypothesis in the context of spatial decision making, we unilaterally photoactivated GABAergic SC neurons while mice performed a spatial choice task. Briefly, the task required mice to sample a binary odor mixture that cues reward location (left or right), wait for a go signal, and execute an orienting movement towards the selected location to retrieve a water reward; light was delivered on 30% of trials while mice selected the reward location^18^ (Fig. 1c; Methods). Given that SC activity represents contralateral targets^8, 11, 41^, including in mice performing this task^13^, we predicted that activating GABAergic SC neurons would inhibit nearby neurons representing the contralateral choice and therefore bias choices ipsilaterally (Fig. 1a,d).

**Figure 1:**
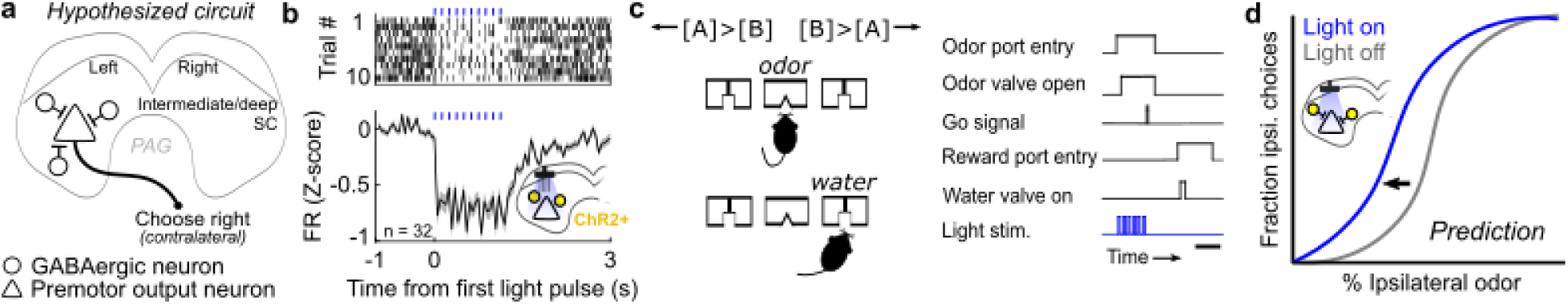
Hypothesis, experiment and predicted results. (**a**) We hypothesized that GABAergic SC neurons locally inhibit premotor output neurons that promote contralateral choices. External inputs to the SC are omitted for simplicity. (**b**) A subpopulation of SC neurons is inhibited by photoactivation of nearby GABAergic neurons, consistent with hypothesis. Top: Raster for example neuron. Blue ticks, light delivery. Bottom: mean z-scored firing rate across neurons inhibited by light (n = 32/301 neurons; p < 0.05, one-tailed Wilcoxon signed-rank test comparing firing rate during light delivery and inter-trial interval). Gray shading, SEM. Here, and in all panels, inset schematic shows experimental setup. (**c**) Spatial choice task. Left: reward location (left or right) was determined by the relative concentrations of Odors A and B [(+)-carvone and (-)-carvone, respectively]. Right: timing of trial events. The mouse enters the central odor port, samples an odor mixture, chooses a reward port, waits for the go signal, executes its choice, and receives water if correct. To test our hypothesis, GABAergic SC neurons were unilaterally photoactivated during spatial choice on 30% of trials. Scale bar, ∼500 ms. (**d**) We predicted that unilateral photoactivation would bias choices ipsilateral to the photoactivated side, indicated by a leftward shift in the psychometric function.

However, inconsistent with our hypothesis, GABAergic SC photoactivation during spatial choice resulted in a significant contralateral choice bias (Fig 2a; n = 96 stimulation/behavior sessions; Supplementary Table 1) compared to the behavior of control Gad2-Cre mice expressing only YFP (Fig 2b; n = 79 sessions; *p* = 2. 4 × 10^-5^, *U* = 5542, two-tailed Mann-Whitney U test). These results suggest one of two possibilities: either local inhibition does not ipsilaterally bias choices in freely orienting mice, or the primary function of GABAergic SC neurons during spatial choice is not to provide local inhibition to premotor output neurons.

**Figure 2:**
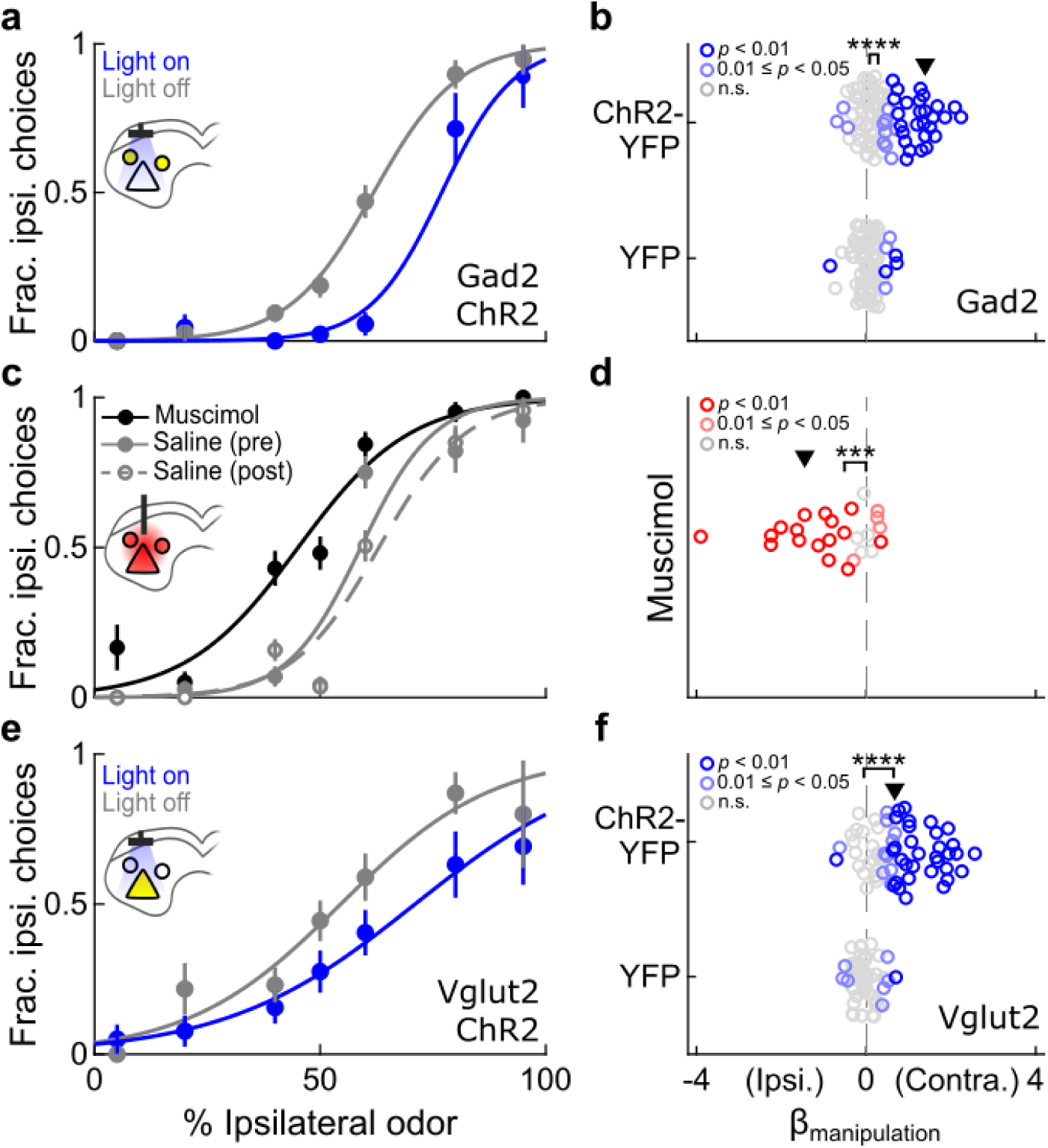
Effects of unilaterally manipulating activity of specific SC cell types on spatial choice. (**a**) Example session in which GABAergic neurons were unilaterally photoactivated during spatial choice. GABAergic activation produced a contralateral choice bias. (**b**) Overall influence of photoactivation on choice in GABAergic (ChR2-YFP; n = 96) and control (YFP; n = 79) sessions, quantified by logistic regression (Methods). Contralateral bias was greater in GABAergic photoactivation than control sessions (**** *p* = 2.4 × 10^-5^, *U* = 5542, two-tailed Mann-Whitney U test). Bracket, difference between group medians (ChR2-YFP = 0.28; YFP = 0.08). ▾, example session in **a**. (**c**) Example sessions in which muscimol or saline was unilaterally delivered to the SC prior to behavioral session. Muscimol produced an ipsilateral choice bias. (**d**) Overall influence of local inhibition on choice in muscimol sessions (n = 25), quantified by logistic regression. Muscimol biased choices ipsilateral relative to saline sessions (*** *p* = 8.9 × 10^-4^, W = 39, two-tailed Wilcoxon signed-rank test). Bracket, group median (= −0.51) from zero. ▾, example sessions in **c**. (**e**) As in **a**, in which glutamatergic neurons were unilaterally photoactivated. Glutamatergic activation produced a contralateral choice bias. (**f**) As in **b**, for glutamatergic photoactivation (ChR2-YFP; n = 64) and control (YFP; n = 52) sessions. Contralateral bias was greater in glutamatergic photoactivation than control sessions (**** *p* = 7.1 × 10^-10^, *U* = 1931, two-tailed Mann-Whitney U test). Bracket, difference between group medians (ChR2-YFP = 0.65; YFP = −0.05). ▾, example session in **e**.

We performed two sets of complimentary experiments to distinguish between these possibilities. First, we directly tested the effect of local inhibition on spatial choice via unilateral infusion of muscimol, a GABA-A receptor agonist, to the SC of behaving mice (Methods). As expected, muscimol reliably biased choices ipsilaterally (Fig. 2c,d; n = 25 sessions, *p* = 8.9 × 10^-4^, W = 39, two-tailed Wilcoxon signed-rank test), confirming that the effects of local inhibition in freely orienting mice are consistent with previous results in rats^19, 42^ and primates^16, 43, 44^ performing analogous spatial choice tasks. Next, we revisited the premise that glutamatergic SC activity, which neighboring GABAergic neurons presumably inhibit, promotes contralateral choices. We photoactivated glutamatergic SC neurons during spatial choice by restricting ChR2 expression to these neurons in Vglut2-Cre mice (Methods). Again, as expected, we found that choices were biased contralaterally compared to Vglut2-Cre mice expressing only YFP (Fig. 2e,f; ChR2-YFP, n = 64 sessions; YFP, n = 52 sessions, *p* = 7.1 × 10^-10,^ *U* = 1931, two-tailed Mann-Whitney U test), and observed a concomitant decrease in reaction time for contralateral choices and an increase for ipsilateral choices (Supplementary Fig. 2). These unilateral pharmacological inactivation (Fig. 2d) and optogenetic glutamatergic activation (Fig. 2f) results are consistent with a conserved role for the mouse SC in mediating contralateral choices via excitatory output and confirm that local inhibition of this output would result in an ipsilateral choice bias. Given that activating GABAergic SC neurons produces a contralateral choice bias (Fig. 2a,b), our results support the conclusion that, contrary to our hypothesis and to the common motif of local inhibition in other microcircuits^40^, GABAergic SC neurons do not function to locally inhibit premotor output during spatial choice.

### Endogenous GABAergic SC activity corresponds to contralateral choices

The unexpectedness of the contralateral choice bias induced by GABAergic SC photoactivation (Fig. 2a,b) prompted us to examine endogenous GABAergic activity during spatial choice, and to measure how SC activity was affected by our stimulation paradigm. To do so we employed an optogenetic identification strategy (i.e., “optotagging”) to target extracellular recordings to GABAergic SC neurons using the same ChR2-expression strategy described for our stimulation/behavior experiments (Supplementary Fig. 3; Methods). GABAergic neurons were identified as having a short-latency response to light during stimulation/recording sessions (reliably spiking within 5 ms following each light pulse) and a high waveform correlation between light-driven and spontaneous action potentials (r^2^ > 0.95; Supplementary Fig. 3; Methods).

We first confirmed the effect of photoactivating GABAergic SC neurons during spatial choice by applying our optotagging strategy to a subset of stimulation/behavior sessions in which we also recorded neural activity (Fig. 3; n = 15 stimulation/behavior/recording sessions; Supplementary Table 1). We compared single unit firing rates between light-on and light-off trials during the choice epoch (defined as the time from 100 ms after odor valve open until the go signal; Fig. 3a) and found that GABAergic activity increased beyond endogenous levels (Fig. 3b; Ipsi. choices: n = 19 neurons; *p* = 0.0013, two-tailed Wilcoxon signed-rank test; Contra. choices: n = 23 neurons; *p* = 0.0014, two-tailed Wilcoxon signed-rank test). Conversely, light had no overall effect on the activity of unidentified neurons (Fig. 3b; Ipsi. choices: n = 30 neurons; *p* = 0.21, two-tailed Wilcoxon signed-rank test; Contra. choices: n = 42 neurons; *p* = 0.25; two-tailed Wilcoxon signed-rank test), although a small subset of neurons exhibited inhibition (data not shown; Ipsi. choices: n = 3/30 neurons; Contra. choices: n = 8/42 neurons; *p* < 0.05; Mann-Whitney U test), consistent with our stimulation/recording results (Fig. 1b). We next examined whether the contralateral choice bias we observed following GABAergic activation (Fig. 2b) could have resulted from excitatory rebound (a burst of activity following the release from prolonged light-induced inhibition). We therefore analyzed activity immediately following the light delivery period (“post-choice epoch”; Fig. 3a) and found no difference between light-on and light-off trials for either our unidentified (Fig. 3c; Ipsi. choices: n = 30 neurons; *p* = 0.28, two-tailed Wilcoxon signed-rank test; Contra. choices: n = 42 neurons; *p* = 0.92; two-tailed Wilcoxon signed-rank test) or GABAergic (Fig. 3c; Ipsi. choices: n = 19 neurons; *p* = 0.33, two-tailed Wilcoxon signed-rank test; Contra. choices: n = 23 neurons; *p* = 0.16, two-tailed Wilcoxon signed-rank test) populations, suggesting the absence of any rebound effect. This analysis of stimulation/behavior/recording sessions show that light delivery increased activity only in GABAergic neurons and only during the choice epoch, confirming the relationship between increased GABAergic SC activity and contralateral choice suggested by our stimulation/behavior sessions (Fig. 2a,b).

**Figure 3:**
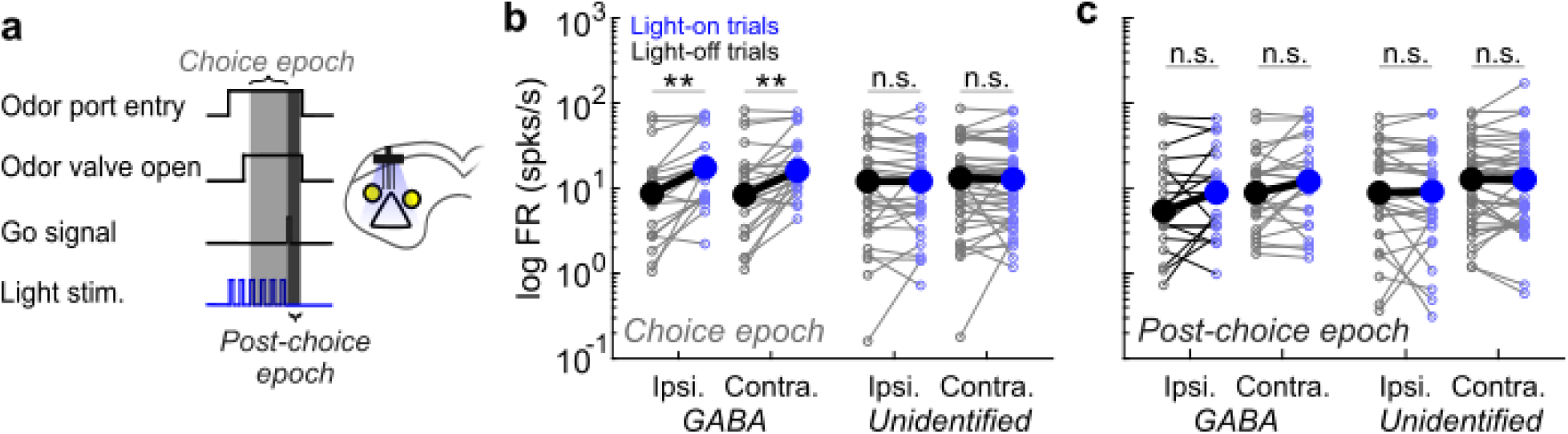
Effects of photoactivation are restricted to GABAergic SC neurons during the choice epoch. (**a**) Schematic of experimental paradigm and epoch definitions, in stimulation/behavior/recording sessions. (**b**) Effect of photoactivation on activity during choice epoch. Activity of GABAergic, but not unidentified, neurons increases on light-on trials compared to light-off trials (GABAergic: Ipsi. choices, n = 19, ** *p* = 0.0013, two-tailed Wilcoxon signed-rank test; Contra. choices, n = 23, ** *p* = 0.0014, two-tailed Wilcoxon signed-rank test; Unidentified: Ipsi. choices, n = 30, *p* = 0.21, two-tailed Wilcoxon signed-rank test; Contra. choices, n = 42, *p* = 0.25, two-tailed Wilcoxon signed-rank test; neurons were excluded from epoch analysis if median firing rate was 0 for the given epoch and direction). Each neuron is represented by a connected pair of small symbols. Large symbols show medians. (**c**) As in **b**, for activity during post-choice epoch. Activity of neither population changes on light-on trials compared to light-off trials (GABAergic: Ipsi. choices: n = 19, *p* = 0.33, two-tailed Wilcoxon signed-rank test; Contra. choices: n = 23, *p* = 0.16, two-tailed Wilcoxon signed-rank test; Unidentified: Ipsi. choices: n = 30, *p* = 0.28, two-tailed Wilcoxon signed-rank test; Contra. choices: n = 42, *p* = 0.92, two-tailed Wilcoxon signed-rank test).

We next examined endogenous activity of GABAergic SC neurons during spatial choice. We sought to determine if activity correlates with contralateral choices, which would be consistent with our manipulation results, or correlates with ipsilateral choices, which would be consistent with our initial hypothesis that GABAergic SC neurons provide local inhibition. By performing our optotagging analysis in conjunction with each behavior/recording session (Supplementary Table 1), we identified 74 GABAergic neurons from a total recorded population of 239 SC neurons that exhibited sufficient activity during the choice epoch (Supplementary Fig 3; Methods), and examined choice-epoch activity separately during ipsilateral and contralateral choices. Figure 4a highlights an example GABAergic neuron showing a greater increase in firing rate during contralateral than ipsilateral choices. We quantified the dependence of firing rate on choice for all recorded neurons by calculating the “choice preference” of each neuron during the choice epoch (Methods). Choice preference is a measure adapted from a ROC-based analysis which uses a sliding criterion to define a curve separating firing rates associated with ipsilateral and contralateral choices (Fig. 4b). The resulting value ranges from −1 (greater activity before ipsilateral choices) to 1 (greater activity before contralateral choices), with 0 indicating no detectable difference in activity. Across all SC neurons with a significant choice preference, more exhibited a contralateral than ipsilateral preference (Fig. 4c, Top; Contra. neurons: 49/72; Ipsi. neurons = 23/72; *p* = 0.0022, Χ^2^ test), consistent with previous results^11–13^. Within the population of GABAergic neurons, we also found that more neurons preferred the contralateral than the ipsilateral choice (Fig. 4c, Bottom; Contra. neurons = 25/29, Ipsi. neurons = 4/29; *p* = 9.7×10^-5^, Χ^2^ test). Consistent with this finding, when we constructed psychometric functions conditional on the choice-epoch firing rate of GABAergic contralateral-preferring neurons, we found that contralateral choices were more likely on trials with higher firing rates (Fig. 4d). Finally, we observed diversity in the timing of peak activity across the population of contralateral-preferring GABAergic neurons (Fig. 4e), with some exhibiting maximal activity early during the choice epoch and others just before the movement was permitted. These results demonstrate that the activity of GABAergic SC neurons reflects contralateral choices, consistent with the contralateral bias elicited by activating these neurons during the choice epoch.

**Figure 4:**
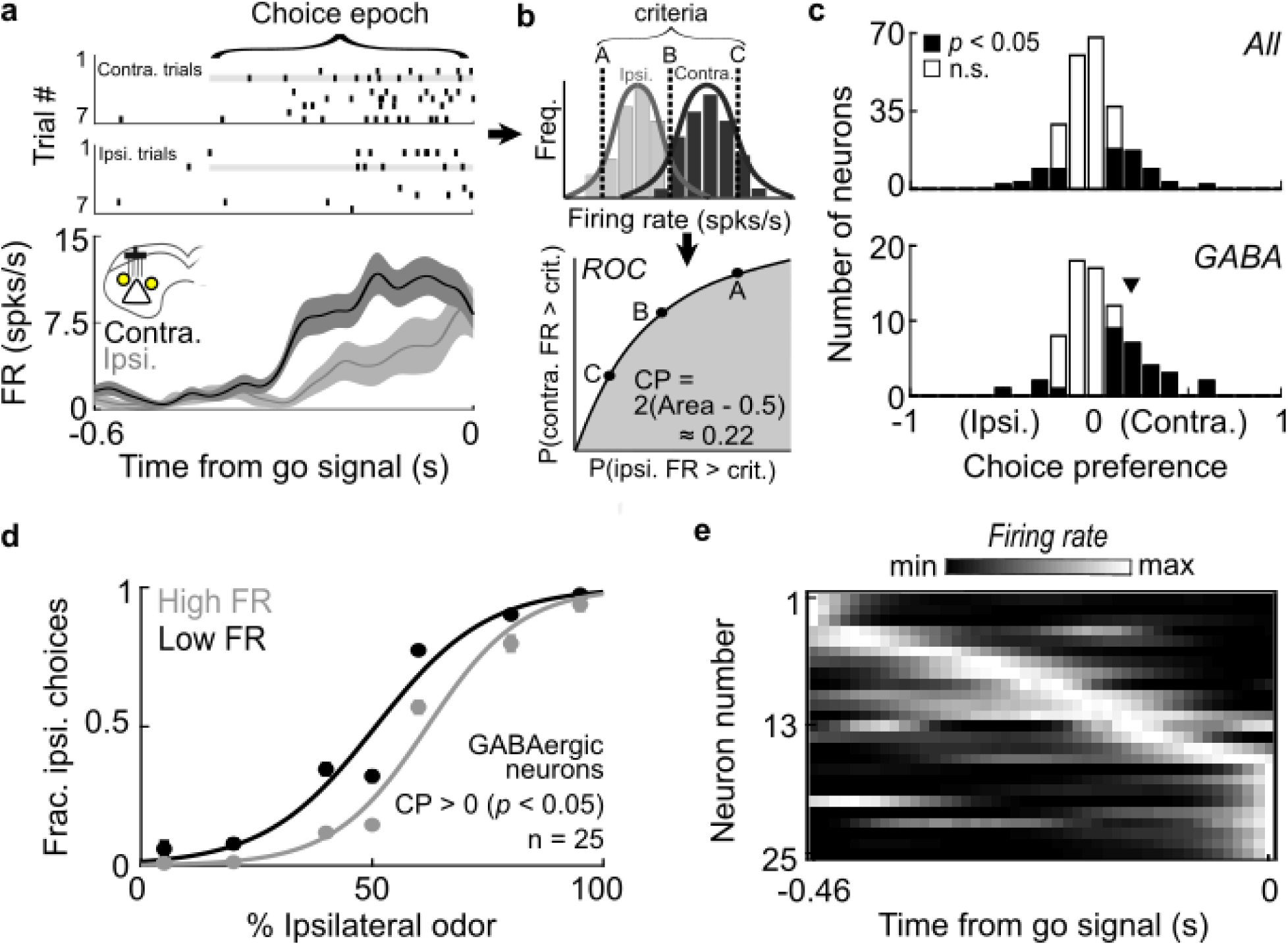
Endogenous activity of GABAergic SC neurons during spatial choice. (**a**) Rasters (top) and peri-event histograms (bottom) for an identified GABAergic neuron (Supplementary Fig. 3) aligned to go signal and segregated by choice. Rasters show 7 randomly selected trials per group. Shading, ± SEM. (**b**) Calculation of choice preference. Choice-epoch firing rate is calculated for each trial (top) and a ROC curve (bottom) is constructed by computing the fraction of trials, separately for ipsilateral and contralateral choices, with higher firing rate than a shifted criterion (e.g., A, B, C). CP = −1 indicates strongest ipsilateral preference and CP = 1 indicates strongest contralateral preference. CP, choice preference. (**c**) Choice preferences during choice epoch for all neurons (top) and GABAergic neurons (bottom). Black, neurons with significant preference (*p* < 0.05). More neurons significantly prefer contralateral than ipsilateral choice (all neurons: 49 prefer contra, 23 prefer ipsi, *p* = 0.0022, Χ^2^ test; GABAergic neurons: 25 prefer contra, 4 prefer ipsi, *p* = 9.7×10^-5^, Χ^2^ test). ▾, example neuron in **a**. (**d**) Conditional psychometric functions constructed separately for trials in which choice-epoch firing rate was high (top quartile) or low (bottom quartile) for each GABAergic neuron with a significant contralateral preference (right black bars at bottom of **c**). Contralateral choice was more likely when firing rate was high. (**e**) Normalized choice-epoch activity on contralateral trials for each GABAergic neuron with a significant contralateral preference (right black bars at bottom of **c**). Neurons are sorted by time of maximum firing rate.

Having established a causal relationship between the activity of GABAergic SC neurons and contralateral choices (Figs. 2, 4), we next utilized our set of stimulation/behavior/recording sessions to examine how tightly GABAergic SC activity and spatial choices are coupled. Specifically, we leveraged the fact that photoactivation elicits a variable number of spikes to test whether the likelihood of contralateral choice on a given trial depended on the number of spikes added on that trial (Fig. 5a). We reasoned that if GABAergic activity makes contralateral choices more likely, then contralateral choices would be more likely to occur on trials with more light-elicited spikes—particularly on “difficult” trials in which the sensory evidence indicating reward side was weak^9,16, 20, 43^. Figure 5b shows, for an example GABAergic neuron, that contralateral choices are more likely on trials in which photoactivation induced more spikes, as demonstrated by a positive exponential fit to trial-by-trial data (Methods). We performed this analysis for each contralateral-preferring GABAergic neuron recorded in stimulation/behavior/recording sessions (n = 5), separately for difficult (% Ipsi. odor = 40, 50, or 60) and easy (% Ipsi. Odor = 5, 20, 80, or 95) trials. We indeed found a stronger relationship between added spikes and choice on difficult than on easy trials (Fig. 5c; *p* = 0.0268, two-tailed paired t-test), further supporting the functional role of GABAergic SC activity in promoting contralateral choice.

**Figure 5:**
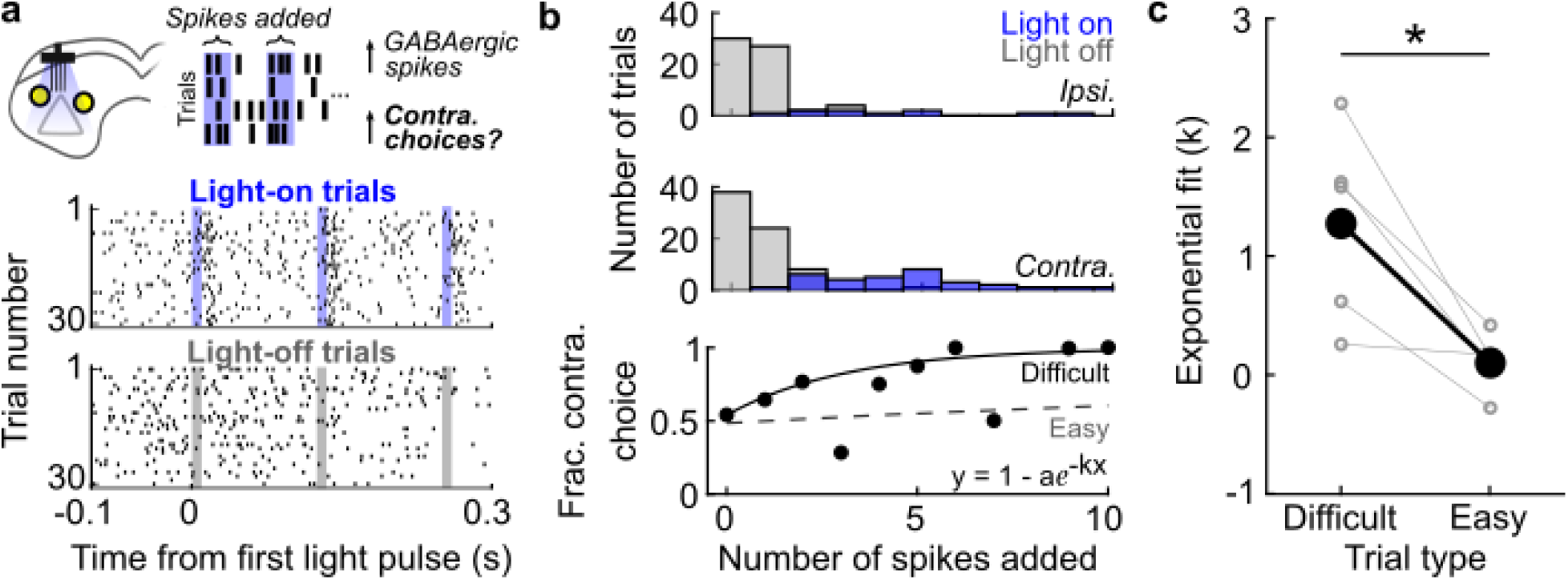
Contralateral choice is more likely on trials in which more spikes in GABAergic SC neurons are elicited by light. (**a**) Top: Schematic of experimental set-up. Rasters for an example contralateral preferring GABAergic neuron on contralateral light on (middle) and light off (bottom) trials, aligned to first light pulse. Shading indicates timing of light pulses on light-on trials. (**b**) Number of ipsilateral (top) and contralateral (middle) choices as a function of number of spikes during time of light delivery on light-on and light-off trials. Bottom: Fraction of contralateral choices for each number of added spikes (circles). Solid line, best-fit exponential function to raw trial-by-trial values (not shown; ipsilateral choice = 0; contralateral choice = 1) for difficult trials (% Ipsi. odor = 40, 50, or 60). Dashed line, same fit for easy trials (% Ipsi. odor = 5, 20, 80, or 95). (**c**) For each GABAergic neuron with a significant contralateral choice preference recorded in stimulation/behavior/recording sessions (n = 5), slope of exponential fit (*k*) is plotted for difficult and easy trials. Slope is larger for difficult than easy trials (* *p* = 0.027, two-tailed paired t-test).

### Intercollicular GABAergic SC neurons mediate spatial choice

Together, the results of our manipulation and recording experiments support an alternative hypothesis: that intrinsic SC inhibition contributes to spatial decision making via non-local mechanisms. To gain insight into these mechanisms we utilized a rate-based bump attractor model^13^ (Methods), similar to those used to elucidate other functions of the SC such as working memory^15^ and countermanding behavior^45^.

In our model the population of excitatory (*E*) and inhibitory (*I*) cells (in a 2:1 ratio) in each colliculus (Fig. 6a) are interconnected using specific intracollicular (intra-SC) distance-dependent rules, where *I* cells can influence neural activity at overall greater distances than *E* cells. A small subpopulation of *E* and *I* cells also form sparse intercollicular (inter-SC) connections to mimic experimental observations^38, 46^.

**Figure 6:**
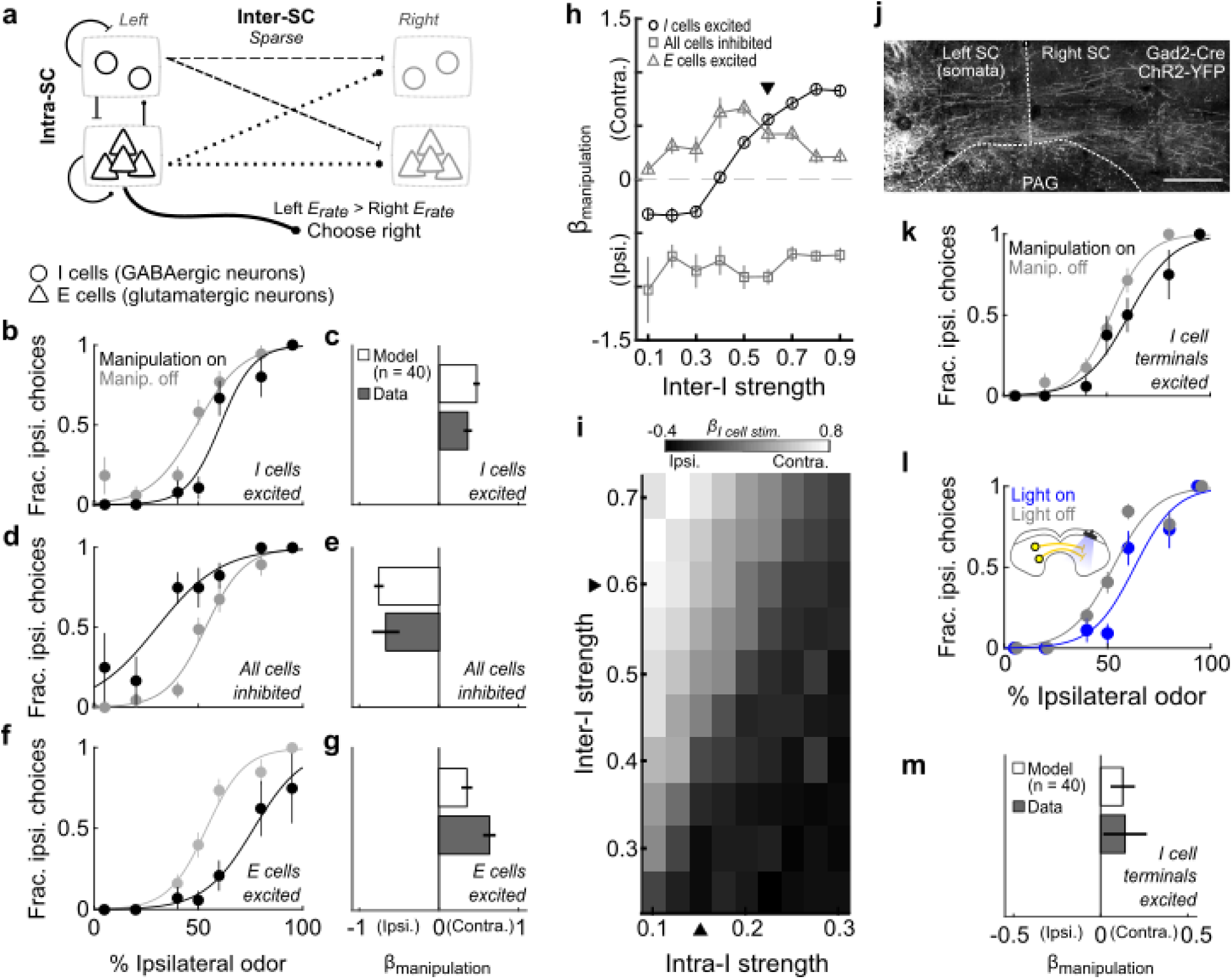
Attractor model recapitulates results when intercollicular inhibition is stronger than intracollicular inhibition. (**a**) Schematic of attractor connectivity. (**b**) Unilateral excitation of inhibitory (*I*) cells during spatial choice in the model (compare to Fig. 2a). (**c**) Influence of unilaterally exciting *I* cells in model sessions compared to experimental photoactivation of GABAergic neurons (data in Fig. 2b). (**d**) Unilateral inhibition of *E* and *I* cells in the model (compare to Fig. 2c). (**e**) Influence of unilaterally inhibiting *E* and *I* cells in model sessions compared to experimental local inhibition (data in Fig. 2d). (**f**) Unilateral excitation of *E* cells during spatial choice in the model (compare to Fig. 2e). (**g**) Influence of unilaterally exciting *E* cells in model sessions compared to experimental photoactivation of glutamatergic neurons (data in Fig. 2f). (**h**) Effect of manipulation in model sessions depends on strength of intercollicular projections of *I* cells (inter-*I*). Each symbol shows mean ± SEM. ▾, strength in **b-g**. (**i**) Effect of exciting *I* cells depends on relative strength of their intercollicular (inter-*I*) and intracollicular (intra-*I*) projections. Mean of 40 sessions per square. ▴, strengths in **b-g**. (**j**) Coronal histological section showing intercollicular GABAergic projections. Scale bar, 100 µm; PAG, periaqueductal gray. (**k**) Excitation of *I* cell intercollicular terminals during spatial choice in the model. (**l**) Stimulation/behavior session in which the intercollicular terminals of GABAergic neurons were photoactivated during spatial choice. (**m**) Influence of exciting intercollicular terminals of *I* cells in model sessions compared to experimental photoactivation of intercollicular terminals of GABAergic neurons (n = 13 sessions).

We initially selected for model parameters for which a sustained locus of activity emerged following external stimulation of *E* and *I* cells. On each model trial, we read out the choice as the side contralateral to the location of highest mean *E* cell activity (left or right SC). This model allowed us to test, over a range of valid parameters, the effect of manipulating the activity of specific cell types, and thereby determine which architectures were consistent with our experimental data.

To model photoactivation of GABAergic neurons (Fig. 2a,b), we unilaterally increased the rate of a subset of *I* cells on 30% of trials during a time period corresponding to the choice epoch in each session (Methods). We found that this manipulation reproduced the contralateral bias observed in our experiments (Fig. 6b,c). Using the same parameters, we modeled pharmacological SC inhibition (Fig. 2c,d) by unilaterally decreasing the activity of a subset of *E* and *I* cells and photoactivation of glutamatergic SC neurons (Fig. 2e,f) by unilaterally increasing the rate of a subset of *E* cells on 30% of trials in each session. For both sets of experiments, we again found that the model frequently reproduced the choice bias we observed experimentally (Fig. 6d-g). We next examined how these results depended on specific model parameters and found that the effect of exciting *I* cells (Fig. 6b,c) strongly depended on the strength of the intercollicular inhibitory (inter-*I*) connections: low values of this parameter yielded an ipsilateral bias, while high values yielded a contralateral bias (Fig. 6h). We further found that the absolute inter-*I* strength did not entirely determine the direction of the choice bias under *I* cell excitation; rather the ratio of inter-*I* to intracollicular inhibition (intra-*I*) strength was critical: exciting *I* cells resulted in the experimentally observed contralateral choice bias when the inter-*I* strength was at least 2.5 times higher than the intra-*I* strength. These modeling results suggest that intercollicular inhibition is a mechanism by which GABAergic SC neurons promote contralateral choice (Figs. 2a,b; 4; 5).

Moreover, the model generated an interesting and testable prediction: activating the terminals of GABAergic neurons in the opposite SC would result in a similar contralateral choice bias. Indeed, we observe intercollicular GABAergic projections in our mice (Fig. 6j) and, when using the same set of parameters as in our other modeling sessions (Figs. 6b-g), we found that unilaterally exciting the terminals of a subset of *I* cells elicited a contralateral bias (Fig. 6k). We then tested this prediction experimentally by unilaterally photoactivating the terminals of GABAergic SC neurons in the opposite SC, which we achieved by replicating the photoactivation experiments shown in Fig. 2a,b but with the optical fiber targeted to the opposite SC (Methods). Consistent with the model predictions, we found that terminal photoactivation resulted in a choice bias contralateral to the ChR2-expressing somata (Fig. 6l,m), and a corresponding decrease in reaction time (Supplementary Fig. 4). Together, these modeling and experimental results support a key role for intercollicular inhibition in spatial choice.

## Discussion

Elucidating the roles of specific cell types in decision making is a critical step toward understanding functional neural circuits and biologically plausible theoretical frameworks for decisions. In this study, we leveraged behavioral, optogenetic, electrophysiological and computational approaches to identify the functional role of inhibitory neurons in the SC in spatial choice, an important and tractable form of decision making for which the SC plays a critical role conserved across species^7, 13^. We initially hypothesized that GABAergic SC neurons function to locally inhibit glutamatergic output representing contralateral choices (Fig. 1a) and thus predicted that activating GABAergic SC neurons would bias mice to select the ipsilateral choice (Fig 1d). Surprisingly, unilateral manipulation of, and extracellular recordings from, GABAergic SC neurons revealed that their activity promotes contralateral choices. Examining the behavior of an attractor model based on our experimental results suggested a role for GABAergic SC neurons in intercollicular inhibition (Fig. 6a-i), which were then supported by the results of terminal photoactivation experiments (Fig. 6l,m). These results offer new insight into how the SC mediates spatial choice and significantly extend our understanding of inhibition in decision making circuits^30–32^.

The computations underlying decision making are generally considered within one of two broad frameworks: competitive winner-take-all models, which require long-range inhibitory interactions between loci representing competing choices^32^, and race-to-threshold models, which do not require such inhibitory interactions^28^. By demonstrating a role for intercollicular inhibition, our data lend cautious support to the former class of models. Despite previous physiological evidence for inhibition between the left and right SC^38, 39^, critical evidence argues against the winner-take-all model of spatial choice in the SC. Most importantly, activity representing competing options builds up in multiple left and right SC loci, and even non-selected representations persist even after the decision is made^11, 12, 27^. One study that directly examined activity in SC loci representing competing choices failed to observe decreases in one locus simultaneous with increases in the other, as would be expected if the two loci mutually inhibit one another^27^. However, we might not expect to observe strict winner-take-all mechanisms in the SC, whereby activity in all but the ‘winning’ locus is silenced, since most natural actions require the rapid execution of a sequence of movements that are represented in the activity of SC loci^47^. Instead, intercollicular inhibition may provide a “winner-take-most” mechanism for making one option more likely to be selected by enhancing the contrast in activity between loci without silencing one entirely. Our data are also consistent with a role for GABAergic SC neurons in setting criteria for decisions requiring stimulus detection^21^; future studies can further elucidate their role by examining these, and other, forms of SC-dependent decisions, including approach vs. avoidance behavior^48^.

Despite GABAergic SC neurons exhibiting no difference in activity between easy and difficult trials (Supplementary Fig. 5), activating them had a stronger influence on choice on difficult trials compared to easy trials (Fig. 5c). The psychometric functions plotted conditionally on GABAergic SC activity (Fig. 4d) show a similar relationship: the difference in choice between low and high activity trials is greatest for the more difficult trials (% Ipsi. odor = 40, 50, or 60). This effect could reflect the balance between intrinsic SC processing and influences extrinsic to the SC such as the basal ganglia and cortical regions, among others, that are known to contribute to spatial choice^9^. When the sensory evidence for the rewarded choice is strong (i.e., on easy trials), input from upstream regions may dominate processing in the SC (or outweigh SC contributions at downstream circuits in the brainstem). Conversely, when the sensory evidence is weak (i.e., on difficult trials), intrinsic SC processing, including via the GABAergic SC neurons studied here, may have more weight in determining the output of competing choices and therefore exert a stronger influence over spatial choice^13^.

While the function of some SC neurons have been studied during behavior^23–26^, characterization of GABAergic neurons in the intermediate and deep layers has primarily been limited to anatomy and slice physiology^33, 34, 38, 49^. We have identified a functional role for these neurons in mediating spatial choice via intercollicular inhibition (Fig. 7). While some GABAergic SC neurons project directly to the opposite SC^34, 38^, which provides the simplest mechanism for explaining our results and is supported by our terminal activation experiments (Fig. 6l,m), we cannot rule out the possibility of additional contributions by multi-synaptic, or extra-collicular, pathways, as has been described in the barn owl^30^. Indeed, activating intercollicular GABAergic terminals produced a weaker contralateral choice bias than somatic stimulation (Figs. 2a,b; 6l,m) and elicited a surprising decrease in reaction time for ipsilateral choices (Supplementary Fig. 4). These results suggest additional circuitry beyond our new working model (Fig. 7), which can be examined in future studies by applying a similar approach as used here, including simultaneous cell-type-specific recording in both colliculi. A finer dissection of inhibitory circuits for decision making will be feasible as genetic markers for subtypes of GABAergic SC neurons are identified^49^.

**Figure 7:**
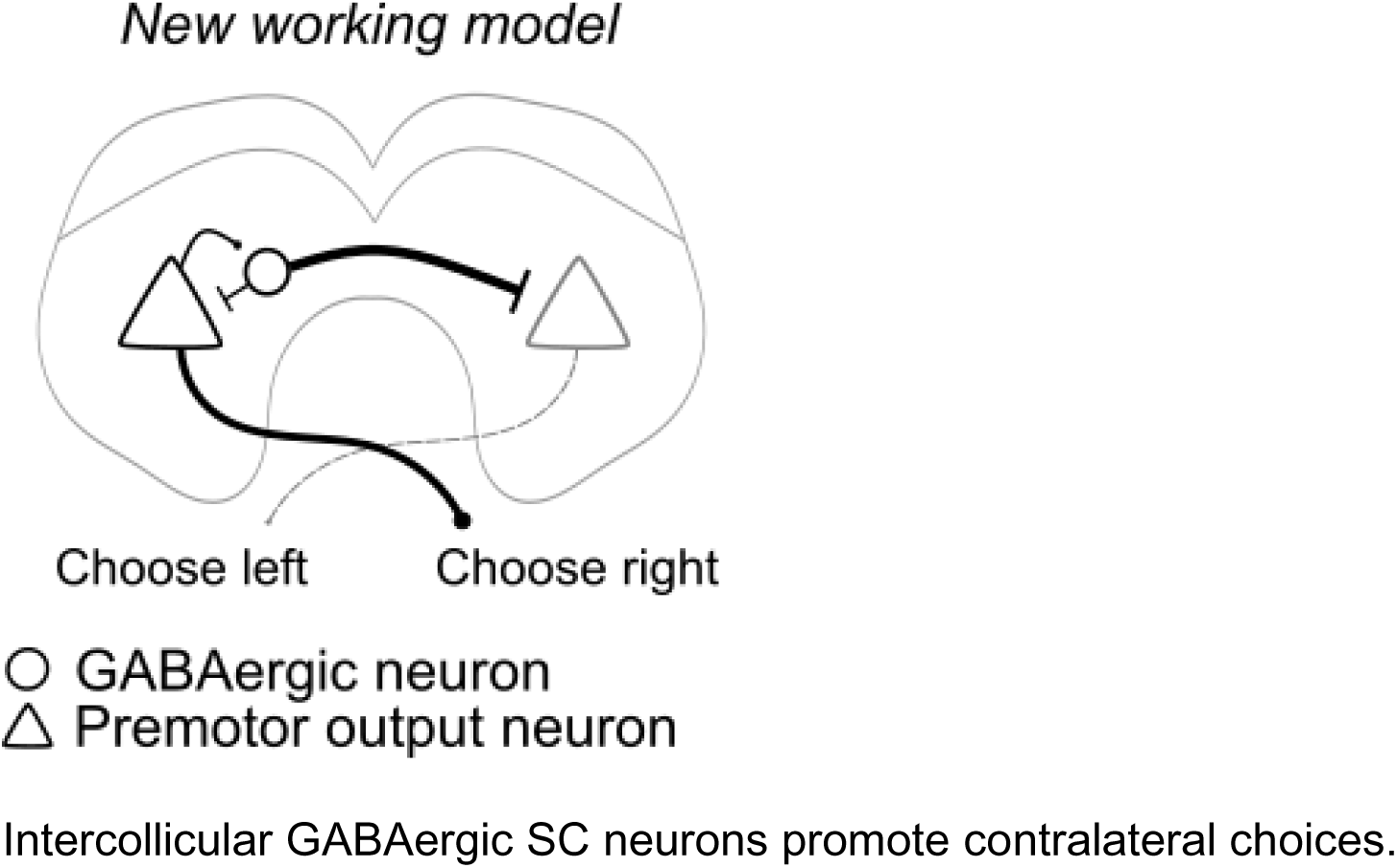
Working model of role of GABAergic SC neurons in spatial choice. Intercollicular GABAergic SC neurons promote contralateral choices.

The motif of long-range inhibition has recently been described in cortical circuits^50^, and is a staple of theoretical models of decision making^30, 32^. Thus, in addition to elucidating the mechanisms of spatial choice in the SC, our findings could inform the function of other neural circuits and provide important constraints on the relative strength of different forms of inhibition in biologically-plausible modeling frameworks.

## Acknowledgements

We thank Drs. Joshua Dudman, Jennifer Hoy, Abigail Person, Dan Tollin, John Thompson and Joel Zylberberg as well as members of the Felsen lab and Matthew Becker for their insightful comments on the manuscript. We thank Nathan D. Baker for technical assistance and Dr. Michael Hall for machining. Light microscopy was performed at the University of Colorado Anschutz Medical Campus Advance Light Microscopy Core and engineering support was provided by the University of Colorado Optogenetics and Neural Engineering Core. Both cores are supported in part by the Rocky Mountain Neurological Disorders Center (P30NS048154), by NIH/NCRR Colorado CTSI grant UL1 RR025780, and by the University of Colorado NeuroTechnology Center. This work was supported by the National Institutes of Health (R01NS079518, F31NS103305).

## Author contributions

J.E and G.F. designed the experiments. J.E. performed recording and behavioral experiments and analyzed the data. J.B.H. performed behavior experiments and analyzed behavioral data. J.E. constructed the model and analyzed the results. J.E. and G.F. guided data analysis and oversaw the project. All authors discussed the results and J.E. and G.F. wrote the manuscript.

## Methods

### Animals

All procedures were approved by University of Colorado School of Medicine Institutional Animal Care and Use Committee. Mice were bred in the animal facilities of the University of Colorado Anschutz Medical Campus or purchased (Jackson Labs). Adult male mice (6-12 months old at time of experiments) from a C57BL/6J background were used in this study. Mice were housed individually in an environmentally controlled room, kept on a 12-hour light/dark cycle and had ad libitum access to food. Mice were water restricted to 1 mL/day and maintained at 80% of their adult weight. Heterozygous Gad2-ires-Cre (Gad2-Cre; Gad2tm2(cre)Zjh/J) mice were used for optogenetic activation and identification of GABAergic neurons (Supplementary Table 1). Homozygous Vglut2-ires-Cre (Vglut2-Cre; Slc17a6tm2(cre)Lowl/J) were used for optogenetic activation of glutamatergic neurons. Muscimol experiments were performed in C57Bl/6 wild-type mice.

### Behavioral task

Mice were trained on a previously published odor-guided spatial-choice task^18, 51^ Briefly, water-restricted mice self-initiated each trial by nose poking into a central port. After a short delay (∼200 ms), a binary odor mixture was delivered. Mice were required to wait 500 ± 55 ms (mean ± SD) for a go signal (a high frequency tone) before exiting the odor port and orienting toward the left or right reward port for water (Fig. 1b). We refer to the time between odor valve open and the go signal as the “choice epoch” (Fig. 3a, 4a). Exiting the odor port prior to the go signal resulted in the unavailability of reward on that trial, although we still analyzed these trials if a reward port was selected. All training and experimental behavioral sessions were conducted during the light cycle.

Odors were comprised of binary mixtures of (+)-carvone (Odor A) and (−)-carvone (Odor B) (Acros), commonly perceived as caraway and spearmint, respectively. In all sessions – including training on the task, as well as during neural recording and manipulation – mixtures in which Odor A > Odor B indicated reward availability at the left port, and Odor B > Odor A indicated reward availability at the right port (Fig. 1c). When Odor A = Odor B, the probability of reward at the left and right ports, independently, was 0.5. The full set of Odor A/Odor B mixtures used was 95/5, 80/20, 60/40, 50/50, 40/60, 20/80, 5/95. Mice completed training in 8-12 weeks and were then implanted with a neural recording drive, optic fiber or drug-delivery cannula as described below. All neural recording and manipulation experiments were performed in mice that were well-trained on the task.

### Stereotactic surgeries

Mice were removed from water restriction and had ad libitum access to water for at least one week before surgery. Preparation for surgery was similar for viral injections and chronic implants. Deep anesthesia was induced with 2% isoflurane (Priamal Healthcare Limited) in a ventilated chamber before being transferred to a stereotaxic frame fitted with a heating pad to maintain body temperature. A nose cone attachment continuously delivered 1.3%–1.6% isoflurane to maintain anesthesia throughout the surgery. Scalp fur was removed using an electric razor and ophthalmic ointment was applied to the eyes. The scalp was cleaned with betadine (Purdue Products) and 70% ethanol before injecting a bolus of topical anesthetic (150 µl 2% lidocaine; Aspen Veterinary Resources) under the scalp. The skull was exposed with a single incision and scalp retraction. The surface of the skull was cleaned with saline and the head was adjusted to ensure lambda was level with bregma (within 150 µm). Immediately following all surgeries, mice were intraperitoneally administered sterile 0.9% saline for rehydration and an analgesic (5 mg/kg Ketofen; Zoetis). A topical antibiotic was applied to the site of incision and mice were given oxygen while waking from anesthesia. Post-operative care, including analgesic and antibiotic administration, continued for up to 5 days after surgery and mice were closely monitored for signs of distress. Additionally, mice recovered after surgery with ad libitum access to water for at least 1 week before being water-restricted for experiments.

For viral injections, a small craniotomy was drilled above the left SC (3.64-4.04 mm posterior to bregma, 0.75-1.25 mm lateral of midline, 0.85-2.17 mm dorsal from the brain surface^52^). To maximize overlap between ChR2 expression and the optetrode in Gad2-Cre mice, up to 3 injections were made within 0.2 mm^2^. Viruses were delivered with a thin glass pipette at an approximate rate of 100–200 nl/min via manual pressure applied to a 30 ml syringe. Pipets remained at depth for 10 min following each injection before retraction from the brain. For optogenetic manipulation and identification of GABAergic neurons, Gad2-Cre (n= 11) mice were injected with a total (across all injections) of 200 nl of DIO-ChR2-eYFP (AAV2.Ef1α.DIO.ChR2.eYFP, UNC Vector Core, 4.2×10^12^ ppm). Vglut2-ires-Cre (n = 4) mice were injected with 200 nl of the same virus to express ChR2 in glutamatergic neurons. Control mice expressing only YFP (Gad2-Cre (n= 2) and Vglut2-Cre (n = 2)) were injected with 100nl DIO-eYFP (AAV2.Ef1α.DIO.eYFP, UNC Vector Core, 4.6×10^12^ ppm). After injection, the skin was sutured (except for the control mice; see below) and mice recovered for 1 week before being water restricted for behavioral training. Expression occurred during the ∼10 weeks of training.

In Vglut2-Cre and control mice, immediately following virus injection, a 105 µm diameter optic fiber (Thorlabs) was implanted in the same craniotomy used for virus injection. The fiber was housed in a ceramic ferrule (Precision Fiber Products MM-FER2007C) and slowly lowered to be slightly (∼200 μm) above the injection site. The fiber was affixed to the skull via a single skull screw, luting (3M), and dental acrylic (A-M Systems).

To deliver light to, and extracellularly record from, the same population of SC neurons, an optetrode drive, an optic fiber surrounded by four tetrodes (see^53^ for construction), was chronically implanted above the DIO-ChR2-eYFP injection site in fully trained Gad2-Cre mice. A large (∼1 mm^2^) craniotomy was made around the initial injection site. Three additional small craniotomies were made anterior to the initial injection site: one for implanting a ground wire and two for skull screws. The drive was slowly lowered into the large craniotomy and secured in place with luting and dental acrylic.

To deliver light to GABAergic terminals in the contralateral (right) SC, a subset (n = 6) of Gad2-Cre mice with optetrode drives were also implanted with an additional 105 µm diameter optic fiber in the contralateral SC. The fiber was implanted at a ∼50° angle from midline to allow for easy access when attaching the patch cable, at the same anterior/posterior location to the optetrode drive, 3 mm lateral of midline and 2.3 mm dorsal from the brain surface. The fiber was fixed to the skull with luting and dental cement.

To infuse muscimol into the intermediate and deep layers of the SC, a steel guide cannula and removable steel insert assembly (Invivo1) was targeted to 0.8 mm dorsal to the surface of the SC in wild-type mice (n = 4). The guide cannula was affixed to the skull with one skull screw, luting and dental acrylic.

### Optogenetic modulation of the SC

A diode-pumped, solid-state laser (473 nm; Shanghai Laser & Optics Century) delivered light through a 105-μm/0.22 numerical aperture patch cable (Thorlabs) was attached to the implanted ferrule or optetrode drive using a ceramic sleeve (Precision Fiber Products SM-CS125S) and index matching solution (Thorlabs G608N3). To activate ChR2+ GABAergic neurons in Gad2-Cre mice during behavior (“stimulation/behavior”; Fig. 2a,b; Supplementary Table 1; n = 96 sessions; 11 mice), power was calibrated with an optic meter before each session (ThorLabs) to deliver a range of laser powers across mice: 58 sessions between 4.4 – 54.8 mW/mm^2^, 23 sessions between 55.9 – 192.3 mW/mm^2^, and 15 sessions between 220 – 427.2 mW/mm^2^ at 8 Hz (10 ms on) (for a small sample (29 sessions), light was delivered at 25 Hz (10 ms on)) during the entire duration of the odor sampling epoch (odor port entry to go signal). The behavioral effects observed did not depend on power or stimulation frequency. Different neural populations were stimulated as the optetrode was advanced through the SC. To activate ChR2+ glutamatergic neurons during the odor sampling epoch in Vglut2-Cre mice (Fig. 2e,f; n = 64 sessions; 4 mice), laser power was delivered at between 3.7 mW/mm^2^ and 29.4 mW/mm^2^ to measure a choice bias without inducing overt orienting movements of the head; these mice were stimulated during the odor sampling epoch at 8 Hz (10 ms on). Gad2-Cre mice expressing only eYFP (Fig. 2b; n = 79; 2 mice) were stimulated at a range of laser powers: 27 sessions at 257.6 mW/mm^2^ and 52 sessions at 441.6 mW/mm^2^ at 8 Hz (10 ms on) during the odor sampling epoch. Vglut2-Cre mice expressing only eYFP (Fig. 2f; n = 52 sessions; 2 mice) were stimulated at a range of laser powers: 30 sessions at 55.2 mW/mm^2^ and 22 sessions at 441.6 mW/mm^2^ at 8 Hz (10 ms on) during the odor sampling epoch. In all Vglut2-Cre and all Gad2-Cre control mice, the same site was stimulated for all sessions. Across all mice and sessions, stimulation occurred randomly on 30% of trials. Extracellular recordings were acquired during 15 photoactivation sessions in Gad2-Cre mice (“stimulation/behavior/recording”; Fig. 3, 5; Supplementary Table 1).

### Muscimol infusion

Prior to each session, an injection cannula was prepared with either muscimol or saline and inserted into the chronically implanted guide cannula while mice were anesthetized. An infusion pump (Harvard Apparatus) was used to administer 300 nL of solution at 0.15 μL/min. Muscimol dosage was 0.1 mg/ml (in saline) and did not induce ipsilateral circling behaviors. After infusion, the internal cannula remained in place for 3 min before it was retracted. Mice recovered from anesthesia in their cage for at least 10 min before beginning the behavioral session.

### Behavioral analysis

The effect of photoactivating GABAergic or glutamatergic SC neurons and locally inhibiting SC neurons with muscimol (“stimulation/behavior”; β_manipulation_; Figs. 2b,d,f, 6c,e,g,m) on choice behavior, was assessed by logistic regression using Python (Sci-Kit Learn package (version 0.21.3)). The logistic function of the form 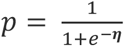, where *p* is the choice made on a given trial (contralateral choice, *p* = 0; ipsilateral choice, p = 1) and *η* is the linear predictor, was adapted to specific analyses as described below. For all analyses, *η* consisted of at least *η*_0_ = *β*_0_ + *β*_*Odor:I*_*x*_*Odor:I*_ + *β*_*Odor:C*_*x*_*Odor:C*_ + *β*_*manipulation*_*x*_*manipulation*_, where *β*_0_ represents overall choice bias, *x*_*Odor:I*_ and *x*_*Odor:C*_ represent the strength of the odors associated with the ipsilateral and contralateral reward port, respectively, *β*_*Odor:I*_ and *β*_*Odor:C*_ represent the influence of the odors on choice, and *β*_*manipulation*_ represents the influence of manipulation (i.e., light or muscimol). *x*_*Odor:I*_ and *x*_*Odor:C*_ are calculated as (fraction of odor - 0.5) / 0.5 and range from 0 to 1; we used separate terms for left and right odors to allow for the possibility that they asymmetrically influenced choice. We set *x*_*manipulation*_ = 0 on light-off /saline trials and *x*_*manipulation*_ = 1 on light-on/muscimol trials. Under these calculations positive β_*manipulation*_ values correspond to ipsilateral influence; for consistency with the choice preference analysis (see below), all β_*manipulation*_ were subsequently multiplied by −1 so that positive values correspond to contralateral influence)^54^. Additionally, L2 regularization (C = 1; default parameter value) was applied to all sessions to account for occasional perfect mouse performance. Sessions were included in analyses only if at least 15 light-on trials and at least 100 total trials (light-on + light-off) were completed. Significance levels for the estimates of β_*manipulation*_ were obtained by shuffling the values of x_*manipulation*_ within a session and recalculating β_*manipulation*_, repeating this process 5000 times to produce a null distribution of β_*manipulation*_ values, and comparing the actual β_*manipulation*_ to the null distribution.

For display, behavioral data from example experimental (Figs. 2a,c,e; 4d) and model (Fig. 6b,d,f,k,l) sessions were fit with the above logistic function with *η* = *β*_0_ + *β*_*Odor*_*x*_*Odor*_, where *x* is the proportion of the ipsilateral odor in the mixture, separately for trials with and without manipulation.

Additionally, we quantified the effects of SC manipulation on reaction time by examining the duration between the go signal and odor port exit. For trials on which mice exited the odor port before the go signal, we calculated the mean time the go signal could have occurred. Significance for each session was calculated using a two-tailed Mann-Whitney U-test to compare reaction times between light-on trials and light-off trials separately for each direction.

### Electrophysiology

Extracellular neuronal recordings were collected using four tetrodes (a single tetrode consisted of four polyimide-coated nichrome wires (Sandvik; single-wire diameter 12.5 μm) gold plated to 0.25-0.3 MΩ impedance). Electrical signals were amplified and recorded using the Digital Lynx S multichannel acquisition system (Neuralynx) in conjunction with Cheetah data acquisition software (Neuralynx).

To sample independent populations of neurons, the tetrodes were advanced between 6 - 23 h before each recording session. To estimate tetrode depths during each session we calculated distance traveled with respect to rotation fraction of the thumb screw of the optetrode drive. One full rotation moved the tetrodes ∼ 450 μm and tetrodes were moved ∼ 100 μm between sessions. The final tetrode location was confirmed through histological assessment using tetrode tracks.

Offline spike sorting and cluster quality analysis was performed using MClust software (MClust 4.4.07, A.D. Redish) in MATLAB (2015a). Briefly, for each tetrode, single units were isolated by manual cluster identification based on spike features derived from sampled waveforms. Identification of single units through examination of spikes in high-dimensional feature space allowed us to refine the delimitation of identified clusters by examining all possible two-dimensional combinations of selected spike features. We used standard spike features for single unit extraction: peak amplitude, energy (square root of the sum of squares of each point in the waveform, divided by the number of samples in the waveform), valley amplitude and time. Spike features were derived separately for individual leads. To assess the quality of identified clusters we calculated isolation distance, a standard quantitative metric^55^. Clusters with an isolation distance > 6 were deemed single units. Units were clustered blind to interspike interval, and only clusters with few interspike intervals < 1 ms were considered for further examination. Furthermore, we excluded the possibility of double counting neurons by ensuring that the waveforms and response properties sufficiently changed across sessions. If they did not, we conservatively assumed that we recorded twice from the same neuron, and only included data from one session.

Electrophysiological recordings were obtained from 308 SC neurons in 96 behavioral sessions from 10 Gad2-Cre mice used in photoactivation experiments. Details of our analyses of the data obtained from our recording experiments are described below. All neural data analyses were performed in MATLAB (2015a/2019a). Neurons recorded during recording/behavior sessions with fewer than 40 trials in either direction or with a choice-epoch firing rate below 2.5 spikes/s for trials in both directions were excluded from all analyses. Additionally, neurons recorded during stimulation/recording/behavior sessions were included in analyses if at least 10 light-on trials were completed in each direction.

### Locally inhibited neuronal responses

To assess local inhibitory effects of GABAergic photoactivation, each neuron recorded during stimulation/recording sessions (n = 301) was tested for a light-induced decrease in firing rate (FR) by comparing the baseline FR (calculated over a 1 s window preceding light delivery [inter-trial interval]) to the FR during photoactivation (entire light delivery period, ∼1.2 s) using a one-tailed Wilcoxon signed-rank test. Neurons with a significant light-induced decrease in FR (*p* < 0.05; n = 32) were Z-score normalized to baseline FRs using 50 ms bins (Fig. 1b).

### Optogenetic identification of GABAergic neurons

Before and/or after behavioral sessions in Gad2-Cre mice expressing ChR2, light was delivered via a diode-pumped, solid-state laser (473 nm; Shanghai Laser & Optics Century) at 8 Hz (10 ms on) or, for a small sample (< 20 sessions), at 25 Hz (10 ms on) to the same population of neurons that were recorded during behavior while also recording extracellular responses to the light (“stimulation/recording”). Isolated units (Supplementary Fig. 3) were identified as GABAergic based on reliable, short-latency responses to light and high waveform correlations between spontaneous and light-evoked action potentials ^56–60^. To correct for cells with high tonic firing rates, responses (i.e., action potentials) had to occur within 5 ms of the onset of light and at a significantly higher probability than responses in 5 ms bins outside of light delivery (i.e., baseline responses). To be maximally conservative, we only considered a neuron to be GABAergic if, out of 5000 baseline responses calculated using randomly selected light-off bins, the neuron responded to light more than every baseline response (Supplementary Fig. 3; p < 0.0002) and exhibited a waveform correlation of at least 0.95 (Supplementary Fig. 3; n = 301 total neurons, 90 identified as GABAergic). Neurons identified as GABAergic could be further identified (i.e., “tracked”) during behavior recordings based on spike features and location (i.e., tetrode number and lead number; Supplementary Fig. 3). Neurons from these recordings were also utilized to detect local inhibition in response to light (Fig. 1b).

GABAergic firing rate increases to photoactivation and absence of excitatory rebound^61^ (Fig. 3b,c) were verified by comparing the firing rate of neurons for light-on and light-off trials separately within a single session. Neurons were included in analyses if their median firing rate was > 0 for the epoch and direction analyzed. For example, a neuron may be excluded from the choice epoch analysis for ipsilateral trials but included in analyses of contralateral trials.

### Waveform analysis

Standard waveform analyses were performed to attempt to identify GABAergic SC neurons based on distinguishing waveform features^62^ (Supplementary Fig. 3). Briefly, the average waveform from each lead was calculated; the lead with the highest peak amplitude was used for analysis. The peak of the waveform was equivalent to the maximum voltage reached during depolarization. Pre- and post-valley were the minimum voltage reached before and after depolarization, respectively.

### Histology

Final tetrode location, fiber and cannula placement, and viral expression was confirmed histologically (Supplementary Fig. 1). Mice were overdosed with an intraperitoneal injection of sodium pentobarbital (100 mg/kg; Sigma Life Science) and transcardially perfused with phosphate buffered saline (PBS) and 4% paraformaldehyde (PFA) in 0.1 M phosphate buffer (PB). After perfusion, brains were submerged in 4% PFA in 0.1 M PB for 24hr for post-fixation and then cryoprotected for at least 12hr immersion in 30% sucrose in 0.1 M PB. On a freezing microtome, the brain was frozen rapidly with dry ice and embedded in 30% sucrose. Serial coronal sections (50 µm) were cut and stored in 0.1M PBS. Sections were stained with 435/455 blue fluorescent Nissl (1:200, NeuroTrace; Invitrogen) to identify cytoarchitectural features of the SC and verify tetrode tracks and implant placement. Images of the SC were captured with a 10× or 20x objective lens, using a 3I Marianis inverted spinning disc confocal microscope (Zeiss) and 3I Slidebook 6.0 software. Images were adjusted in ImageJ to enhance contrast.

### Choice preference

To examine the dependence of the firing rate of individual neurons on choice (Fig. 4c), we used an ROC-based analysis^63^ that quantifies the ability of an ideal observer to classify whether a given spike rate during the choice epoch was recorded in one of two conditions (here, during ipsilateral or contralateral choice). We defined the choice epoch as beginning 100 ms after odor valve and ending with the go signal. We defined “choice preference” as 2(ROCarea − 0.5), a measure ranging from − 1 to 1, where – 1 denotes the strongest possible preference for ipsilateral, 1 denotes the strongest possible preference for contralateral, and 0 denotes no preference^64, 65^. Statistical significance was determined with a permutation test: we recalculated the preference after randomly reassigning all firing rates to either of the two groups arbitrarily, repeated this procedure 500 times to obtain a distribution of values, and calculated the fraction of random values exceeding the actual value. We tested for significance at α = 0.05.

Activity during the choice epoch of contralateral-preferring GABAergic neurons (Fig. 4e) was averaged across contralateral trials in 10 ms bins. Each neuron was rescaled during the choice epoch from 0 (lowest activity; black) to 1 (highest activity; white) and smoothed with a gaussian filter (σ = 60 ms). Example peristimulus time histograms in 4a were smoothed with a Gaussian filter (σ = 18 ms).

Conditional psychometric functions were constructed based on choice-epoch firing rate using the same method described above (Behavioral analysis). Firing rates during the choice epoch of each GABAergic neuron with a significant contralateral preference (Fig. 4c, bottom) were sorted from lowest to highest. Psychometric functions were then constructed based on choices made on trials in which firing rate was high (top quartile) or low (bottom quartile).

Choice as a function of the number of light-induced action potentials from contralateral preferring GABAergic neurons recorded during stimulation/behavior/recording sessions (Fig. 5; Supplementary Table 1) was fit to an exponential function, *y* = 1 − *ae*^−*kx*^, where *y* is the choice on each trial (0 = ipsilateral choice; 1 = contralateral choice), *x* is the number of light-induced action potentials, *k* is the slope of the curve (reported free parameter) and *a* is an additional free parameter to set the intercept.

### Attractor model

To study how SC circuitry underlies spatial choice, we constructed a rate-based bump attractor model^13, 66^ consisting of 200 excitatory (*E*) cells and 100 inhibitory (*I*) cells per SC (600 cells total), an *E:I* ratio consistent with the intermediate and deep layers of the SC^33, 34^. Intra-SC synaptic weights were larger for nearby cells, and smaller for more distant ones, determined by

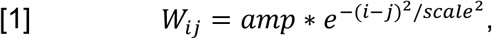

where *i* and *j* are the locations of the pre- and post-synaptic cells, respectively, and *amp* and *scale* are defined independently for presynaptic *E* and *I* cells (*ampE* = 0.01, *ampI *= 0.15, *scaleE *= 0.048, *scaleI *= 0.65). *amp* sets the amplitude (i.e., “strength”) of the connection weights, and *scale* determines the spatial extent over which the connection strength decays. Overall, *I* cells had a higher *scale* than *E* cells, producing conditions for recurrent excitation in the model that are observed experimentally in the SC^67, 68^. Inter-SC synaptic connections were made sparse by assigning synaptic weights to a subset of probabilistically determined contralateral *E* and *I* cells (*InterEamp *= 0.003, *InterIamp *= 0.6, *InterEscale *= 0.3, *InterIscale *= 0.6) while all other weights were set to zero. Inter-SC connections were reciprocated between the left and right SC. To promote network stability, each *W* was normalized to have a maximum eigenvalue of 1.5 by dividing all connection values by max(λ)/1.5, where max(λ) is the largest eigenvalue of the matrix after initialization.

Spike rates of *E* and *I* cells in the left SC (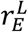 and 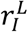, respectively) evolved at each time step (∼ 2 ms in our numerical simulations) according to

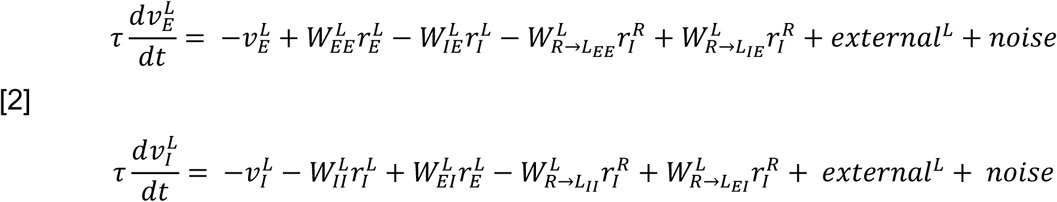

and all spike rates were rectified at each time step according to

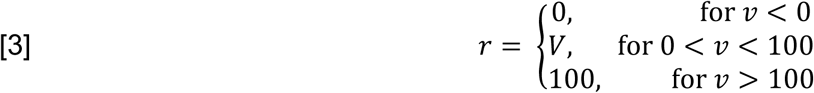

where, e.g., 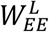 represents synaptic weights from left *E* to *E* cells, 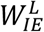 represents weights from left *I* to *E* cellls, and 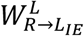represents weights from right *I* to left *E* cells. Noise was drawn from a Gaussian distribution with mean = 0, and variance = 15 for *E* cells and 5 for *I* cells. 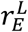 and 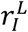 are vectors, with one entry per *E* or *I* cell in the left SC, respectively. Similarly, 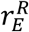 and 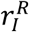 describe the firing rates of cells in the right SC, and they evolve over time via the same equations as those in the left SC (i.e., via Eqs. [2], [3], with all “L”s and “R”s swapped).

The vectors *external*^*L*^ and *external*^*R*^ represent the drive to the *E* and *I* cells from sources outside the SC. For all trial types, external drive was applied in a linearly graded fashion to the last quarter of cells (*E* cells numbered 150 to 200 and *I* cells numbered 75 to 100). On leftward trials, the right SC received a stronger external drive than the left SC, and vice versa in rightward trials. Specifically, on “easy” leftward trials, the drive to left SC cell *i* was *externali^L^* = 0.5i + 0.8, and the drive to right SC cell i was *externali^R^* = 9.5i + 0.8; and on “easy” rightward trials, the drive to left SC cell *i* was *externali^L^* = 9.5i + 0.8, and the drive to right SC cell *i* was *externali^R^* = 0.5i + 0.8. For each trial, external drive began at the start of the “choice epoch” (at time step 151), corresponding to the odor delivery time in the task, and stopped at the end of the choice epoch (at time step 400) for an approximate total time of ∼ 500 ms. At step 401, immediately following the choice epoch, the mean *E* cell firing rate was calculated and choice was determined by the SC with the highest rate. Each “session” consisted of 255 trials, with the proportion of different difficulties matched to the behavioral sessions. Further, to recapitulate mouse-to-mouse experimental variation in viral expression and stimulation efficacy, simulated manipulations occurred at different, randomly selected locations across sessions (although the same cells were manipulated within a single session). To simulate photoactivation of GABAergic neurons, a cluster of 30 *I* cells were given additional excitatory input during the choice epoch by increasing the rate of *I* cells by 50 at 8 Hz (5 steps on). Photoactivation of glutamatergic neurons was simulated by giving addition excitatory input to 30 *E* cells using the same stimulation strategy above, except instead of increasing the rate by 50, the rate was increased by 20. Decreasing the amount of additional excitatory input was analogous to decreasing the laser power for photoactivation experiments (Fig. 2e,f), a tactic employed to reduce overt movements upon Vglut2-ChR2 SC activation (Methods: Optogenetic modulation of the SC).

To simulate muscimol inhibition, a cluster 100 *E* cells and the neighboring 50 *I* cells were given inhibitory input throughout the duration of each trial by subtracting 90 from the rate of these neurons at each step. To simulate inter-SC GABAergic terminal activation, *I* cell terminals were activated on a cluster of 20 *E* cells and 10 *I* cells by giving additional excitatory input, proportional to the synaptic weights of the inter-*I* connections, at 8 Hz during the choice epoch.

All model results were analyzed using the same methods described for behavioral analysis (Methods: Behavioral analysis).

### Statistical analysis

Normal distribution was tested using the Anderson-Darling test and variance was compared. Data were analyzed using MATLAB (2015a/2019a) and presented primarily as medians or means ± SEM. *p* values for each comparison are reported in the figure legends, results and/or methods sections. Mice were excluded from analyses based on the misalignment of the optic fiber with ChR2 expression, lack of ChR2 expression, and misplacement of optetrode outside of the SC (13 mice were excluded from the final data set).

### Data and code availability

The data that support the findings of this study and the custom MATLAB code are available from the corresponding author upon reasonable request.

## References

1. Carandini, M. From circuits to behavior: A bridge too far? Nat. Neurosci. 15, 507– 509 (2012).

2. Shadlen, M. N. & Kiani, R. Decision making as a window on cognition. Neuron 80, 791–806 (2013).

3. Schall, J. D. Accumulators, Neurons, and Response Time. Trends Neurosci. 42, 848–860 (2019).

4. Marr, D. C. & Poggio, T. From understanding computation to understanding neural circuitry. Neurosci. Res. Program Bull. 15, 470–488 (1977).

5. Luo, L., Callaway, E. M. & Svoboda, K. Genetic Dissection of Neural Circuits: A Decade of Progress. Neuron 98, 865 (2018).

6. Krakauer, J. W., Ghazanfar, A. A., Gomez-Marin, A., MacIver, M. A. & Poeppel, D. Neuroscience Needs Behavior: Correcting a Reductionist Bias. Neuron 93, 480–490 (2017).

7. Basso, M. A. & May, P. J. Circuits for Action and Cognition: A View from the Superior Colliculus. Annu. Rev. Vis. Sci. 3, 197–226 (2017).

8. Gandhi, N. J. & Katnani, H. A. Motor functions of the superior colliculus. Annu. Rev. Neurosci. 34, 205–31 (2011).

9. Wolf, A. B. et al. An integrative role for the superior colliculus in selecting targets for movements. J. Neurophysiol. 4532, jn.00262.2015 (2015).

10. Glimcher, P. W. & Sparks, D. L. Movement selection in advance of action in the superior colliculus. Nature 355, 542–5 (1992).

11. Horwitz, G. D. & Newsome, W. T. Target Selection for Saccadic Eye Movements: Prelude Activity in the Superior Colliculus During a Direction-Discrimination Task. J. Neurophysiol. 86, 2543–2558 (2001).

12. Felsen, G. & Mainen, Z. F. Midbrain contributions to sensorimotor decision making. J. Neurophysiol. 108, 135–47 (2012).

13. Lintz, M. J., Essig, J., Zylberberg, J. & Felsen, G. Spatial representations in the superior colliculus are modulated by competition among targets. Neuroscience 408, 191–203 (2019).

14. Steinmetz, N. A., Zatka-Haas, P., Carandini, M. & Harris, K. D. Distributed coding of choice, action and engagement across the mouse brain. Nature 576, 266–273 (2019).

15. Kopec, C. D., Erlich, J. C., Brunton, B. W., Deisseroth, K. & Brody, C. D. Cortical and Subcortical Contributions to Short-Term Memory for Orienting Movements. Neuron 88, 367–377 (2015).

16. McPeek, R. M. & Keller, E. L. Deficits in saccade target selection after inactivation of superior colliculus. Nat. Neurosci. 7, 757–763 (2004).

17. Glimcher, P. W. & Sparks, D. L. Effects of low-frequency stimulation of the superior colliculus on spontaneous and visually guided saccades. J. Neurophysiol. 69, 953–64 (1993).

18. Stubblefield, E. A., Costabile, J. D. & Felsen, G. Optogenetic investigation of the role of the superior colliculus in orienting movements. Behav. Brain Res. 255, 55– 63 (2013).

19. Felsen, G. & Mainen, Z. F. Neural Substrates of Sensory-Guided Locomotor Decisions in the Rat Superior Colliculus. Neuron 60, 137–148 (2008).

20. Wang, L., McAlonan, K., Goldstein, S., Gerfen, C. R. & Krauzlis, R. J. A causal role for mouse superior colliculus in visual perceptual decision-making. Preprint at https://www.biorxiv.org/content/10.1101/835066v1?rss=1 (2019).

21. Crapse, T. B., Lau, H. & Basso, M. A. A Role for the Superior Colliculus in Decision Criteria. Neuron 97, 181–194.e6 (2018).

22. Basso, M. A. & Wurtz, R. H. Modulation of neuronal activity by target uncertainty. Nature 389, 66–69 (1997).

23. Shang, C. et al. A parvalbumin-positive excitatory visual pathway to trigger fear responses in mice. Science 348, 1472–1477 (2015).

24. Evans, D. A. et al. A synaptic threshold mechanism for computing escape decisions. Nature 558, 590–594 (2018).

25. Hoy, J. L., Bishop, H. I. & Niell, C. M. Defined Cell Types in Superior Colliculus Make Distinct Contributions to Prey Capture Behavior in the Mouse. Curr. Biol. 29, 4130–4138.e5 (2019).

26. Zhang, Z. et al. Superior Colliculus GABAergic Neurons Are Essential for Acute Dark Induction of Wakefulness in Mice. Curr. Biol. 29, 637–644.e3 (2019).

27. Ratcliff, R. et al. Inhibition in Superior Colliculus Neurons in a Brightness Discrimination Task? Neural Comput. 23, 1790–1820 (2011).

28. Ratcliff, R., Cherian, A. & Segraves, M. A Comparison of Macaque Behavior and Superior Colliculus Neuronal Activity to Predictions From Models of Two-Choice Decisions. J. Neurophysiol. 90, 1392–1407 (2003).

29. Kim, B. & Basso, M. A. A probabilistic strategy for understanding action selection. J. Neurosci. 30, 2340–55 (2010).

30. Mysore, S. P. & Knudsen, E. I. Reciprocal Inhibition of Inhibition: A Circuit Motif for Flexible Categorization in Stimulus Selection. Neuron 73, 193–205 (2012).

31. Lo, C.-C. & Wang, X.-J. Cortico–basal ganglia circuit mechanism for a decision threshold in reaction time tasks. Nat. Neurosci. 9, 956–963 (2006).

32. Machens, C. K., Romo, R. & Brody, C. D. Flexible control of mutual inhibition: a neural model of two-interval discrimination. Science 307, 1121–4 (2005).

33. Mize, R. R. The organization of GABAergic neurons in the mammalian superior colliculus. Prog. Brain Res. 90, 219–48 (1992).

34. Sooksawate, T., Isa, K., Behan, M., Yanagawa, Y. & Isa, T. Organization of GABAergic inhibition in the motor output layer of the superior colliculus. Eur. J. Neurosci. 33, 421–432 (2011).

35. Hikosaka, O. & Wurtz, R. H. Visual and oculomotor functions of monkey substantia nigra pars reticulata. III. Memory-contingent visual and saccade responses. J. Neurophysiol. 49, 1268–1284 (1983).

36. Lee, P. & Hall, W. C. An in vitro study of horizontal connections in the intermediate layer of the superior colliculus. J. Neurosci. 26, 4763–8 (2006).

37. Phongphanphanee, P. et al. Distinct local circuit properties of the superficial and intermediate layers of the rodent superior colliculus. Eur. J. Neurosci. 40, 2329– 2343 (2014).

38. Takahashi, M., Sugiuchi, Y., Izawa, Y. & Shinoda, Y. Commissural Excitation and Inhibition by the Superior Colliculus in Tectoreticular Neurons Projecting to Omnipause Neuron and Inhibitory Burst Neuron Regions. J. Neurophysiol. 94, 1707–1726 (2005).

39. Munoz, D. P. & Istvan, P. J. Lateral inhibitory interactions in the intermediate layers of the monkey superior colliculus. J. Neurophysiol. 79, 1193–209 (1998).

40. Guo, Z. V et al. Flow of cortical activity underlying a tactile decision in mice. Neuron 81, 179–94 (2014).

41. Wang, L., Liu, M., Segraves, M. A. & Cang, J. Visual Experience Is Required for the Development of Eye Movement Maps in the Mouse Superior Colliculus. J. Neurosci. 35, 12281–6 (2015).

42. Duan, C. A., Erlich, J. C. & Brody, C. D. Requirement of Prefrontal and Midbrain Regions for Rapid Executive Control of Behavior in the Rat. Neuron 86, 1491– 1503 (2015).

43. Lovejoy, L. P. & Krauzlis, R. J. Inactivation of primate superior colliculus impairs covert selection of signals for perceptual judgments. Nat. Neurosci. 13, 261–266 (2010).

44. Nummela, S. U. & Krauzlis, R. J. Inactivation of primate superior colliculus biases target choice for smooth pursuit, saccades, and button press responses. J. Neurophysiol. 104, 1538–48 (2010).

45. Duan, C. A. et al. Collicular circuits for flexible sensorimotor routing. Preprint at https://www.biorxiv.org/content/10.1101/245613v2 (2019).

46. Sooksawate, T., Isa, K., Behan, M., Yanagawa, Y. & Isa, T. Organization of GABAergic inhibition in the motor output layer of the superior colliculus. Eur. J. Neurosci. 33, 421–32 (2011).

47. Port, N. L. & Wurtz, R. H. Sequential Activity of Simultaneously Recorded Neurons in the Superior Colliculus During Curved Saccades. J. Neurophysiol. 90, 1887–1903 (2003).

48. Dean, P., Redgrave, P. & Westby, G. W. Event or emergency? Two response systems in the mammalian superior colliculus. Trends Neurosci. 12, 137–47 (1989).

49. Villalobos, C. A., Wu, Q., Lee, P. H., May, P. J. & Basso, M. A. Parvalbumin and GABA Microcircuits in the Mouse Superior Colliculus. Front. Neural Circuits 12, 35 (2018).

50. Tamamaki, N. & Tomioka, R. Long-Range GABAergic Connections Distributed throughout the Neocortex and their Possible Function. Front. Neurosci. 4, 202 (2010).

51. Uchida, N. & Mainen, Z. F. Speed and accuracy of olfactory discrimination in the rat. Nat. Neurosci. 6, 1224–1229 (2003).

52. Franklin, K. B. J. & Paxinos, G. The mouse brain in stereotaxic coordinates. (Academic Press, Cambridge, 2008).

53. Anikeeva, P. et al. Optetrode: a multichannel readout for optogenetic control in freely moving mice. Nat. Neurosci. 15, 163–170 (2012).

54. Salzman, C. D., Murasugi, C. M., Britten, K. H. & Newsome, W. T. Microstimulation in visual area MT: effects on direction discrimination performance. J. Neurosci. 12, 2331–55 (1992).

55. Schmitzer-Torbert, N., Jackson, J., Henze, D., Harris, K. & Redish, A. D. Quantitative measures of cluster quality for use in extracellular recordings. Neuroscience 131, 1–11 (2005).

56. Cohen, J. Y., Haesler, S., Vong, L., Lowell, B. B. & Uchida, N. Neuron-type-specific signals for reward and punishment in the ventral tegmental area. Nature 482, 85–8 (2012).

57. Cardin, J. A. et al. Targeted optogenetic stimulation and recording of neurons in vivo using cell-type-specific expression of Channelrhodopsin-2. Nat. Protoc. 5, 247–254 (2010).

58. Roux, L., Stark, E., Sjulson, L. & Buzsáki, G. In vivo optogenetic identification and manipulation of GABAergic interneuron subtypes. Curr. Opin. Neurobiol. 26, 88– 95 (2014).

59. Lima, S. Q., Hromádka, T., Znamenskiy, P. & Zador, A. M. PINP: a new method of tagging neuronal populations for identification during in vivo electrophysiological recording. PLoS One 4, e6099 (2009).

60. Pi, H.-J. et al. Cortical interneurons that specialize in disinhibitory control. Nature 503, 521–524 (2013).

61. Li, N. et al. Spatiotemporal constraints on optogenetic inactivation in cortical circuits. Elife 8, (2019).

62. Quirk, M. C., Sosulski, D. L., Feierstein, C. E., Uchida, N. & Mainen, Z. F. A defined network of fast-spiking interneurons in orbitofrontal cortex: responses to behavioral contingencies and ketamine administration. Front. Syst. Neurosci. 3, 13 (2009).

63. Green, D. M. & Swets, J. A. Signal Detection Theory and Psychophysics. (Wiley, New York, 1966).

64. Feierstein, C. E., Quirk, M. C., Uchida, N., Sosulski, D. L. & Mainen, Z. F. Representation of Spatial Goals in Rat Orbitofrontal Cortex. Neuron 51, 495–507 (2006).

65. Crapse, T. B. & Basso, M. A. Insights into decision making using choice probability. J. Neurophysiol. 114, 3039–3049 (2015).

66. Wimmer, K., Nykamp, D. Q., Constantinidis, C. & Compte, A. Bump attractor dynamics in prefrontal cortex explains behavioral precision in spatial working memory. Nat. Neurosci. 17, 431–439 (2014).

67. Pettit, D. L., Helms, M. C., Lee, P., Augustine, G. J. & Hall, W. C. Local Excitatory Circuits in the Intermediate Gray Layer of the Superior Colliculus. J. Neurophysiol. 81, 1424–1427 (1999).

68. Saito, Y. & Isa, T. Local excitatory network and NMDA receptor activation generate a synchronous and bursting command from the superior colliculus. J. Neurosci. 23, 5854–64 (2003).

