## Supplementary Figures and Table for "Inhibitory midbrain neurons mediate decision making"

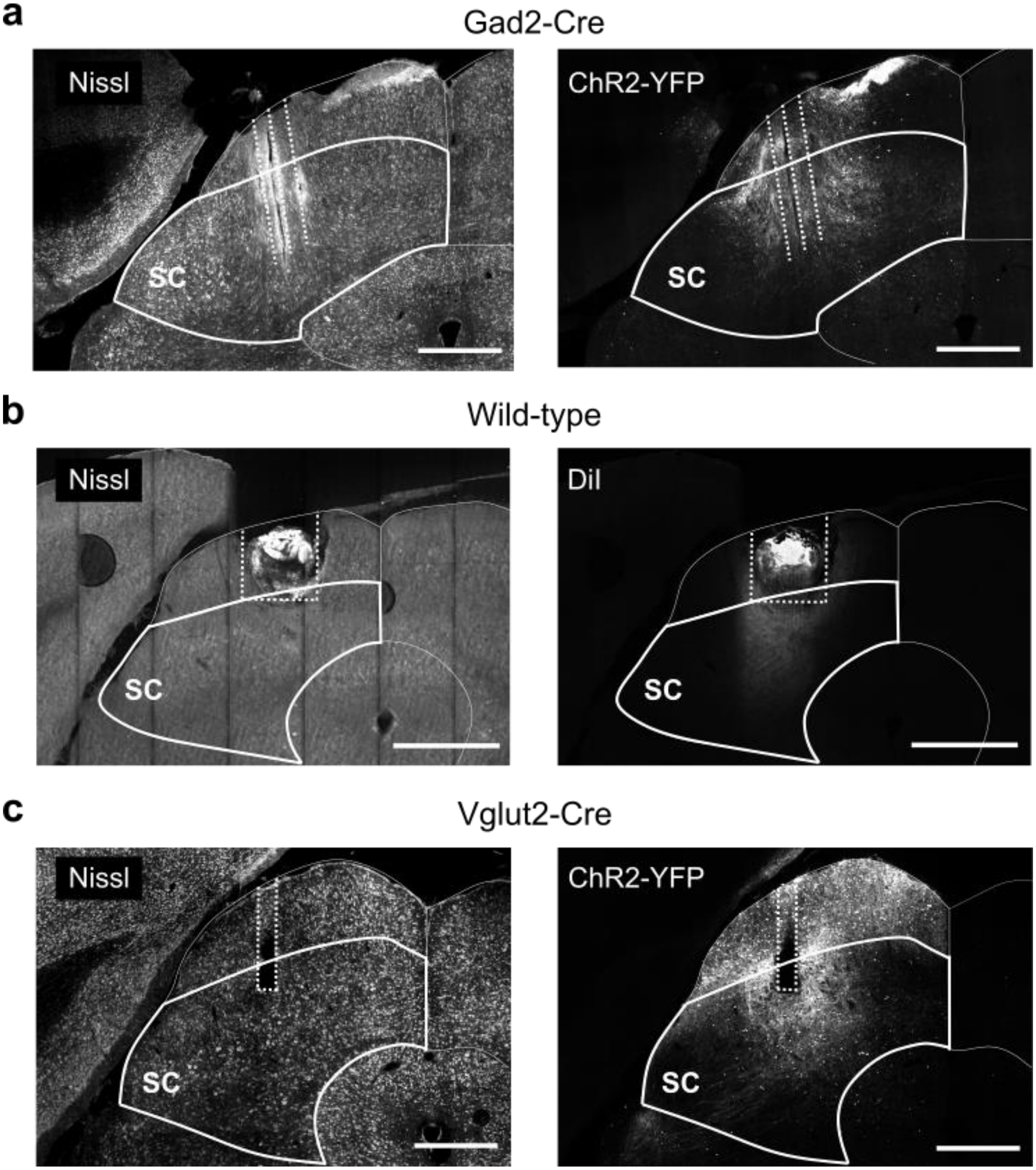


**Supplementary Figure 1: Representative coronal histological sections showing ChR2-YFP expression and targeting of intermediate layer of SC.**
(**a**) Nissl (left) and ChR2-YFP expression (right) in left SC of Gad2-Cre mouse. Dashed lines show visible optetrode tracks. Scale bar, 500 µm. (**b**) Nissl (left) and DiI (right) in left SC of wild-type mouse used in muscimol experiments. DiI coated the internal cannula used for drug delivery. Dashed lines show visible guide cannula track. Scale bar, 500 µm.
(**c**) Nissl (left) and ChR2-YFP expression (right) in left SC of Vglut2-Cre mouse. Dashed lines show visible optical fiber track. Scale bar, 500 µm.

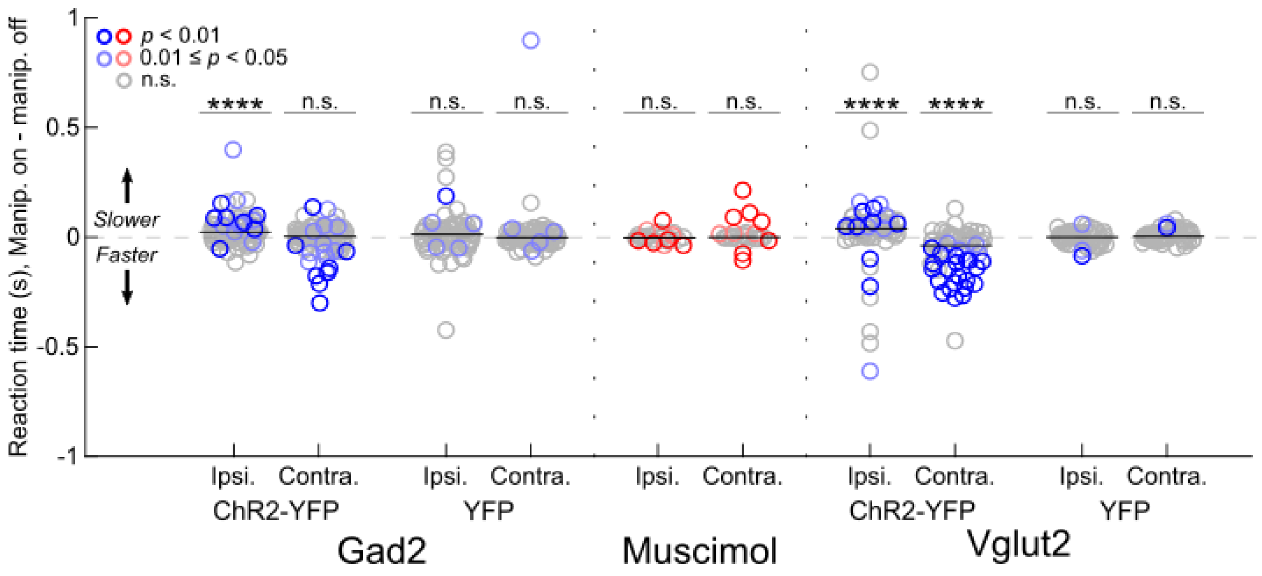


**Supplementary Figure 2: Effect of manipulating SC activity on reaction times.** On each trial, reaction time was calculated as the duration between go signal and odor port exit. In each session, median reaction time was calculated separately for light-on and light-off ipsilateral and contralateral trials, and the difference in medians within each direction is shown, separately for each group of experiments. Each session is represented by 2 symbols (1 for ipsilateral trials and 1 for contralateral trials). Horizontal lines show median within group for each direction. Effect of manipulation is consistent with directional biases shown in Fig. 2 (e.g., when photoactivation biased choices contralaterally, contralateral reaction times tended to be faster on trials with than without photoactivation). For muscimol sessions, reaction times were compared to the preceding and following saline sessions. All groups were compared to zero using a two-tailed Wilcoxon signed-rank test (**** $p < 0.0001$, n.s., not significant).

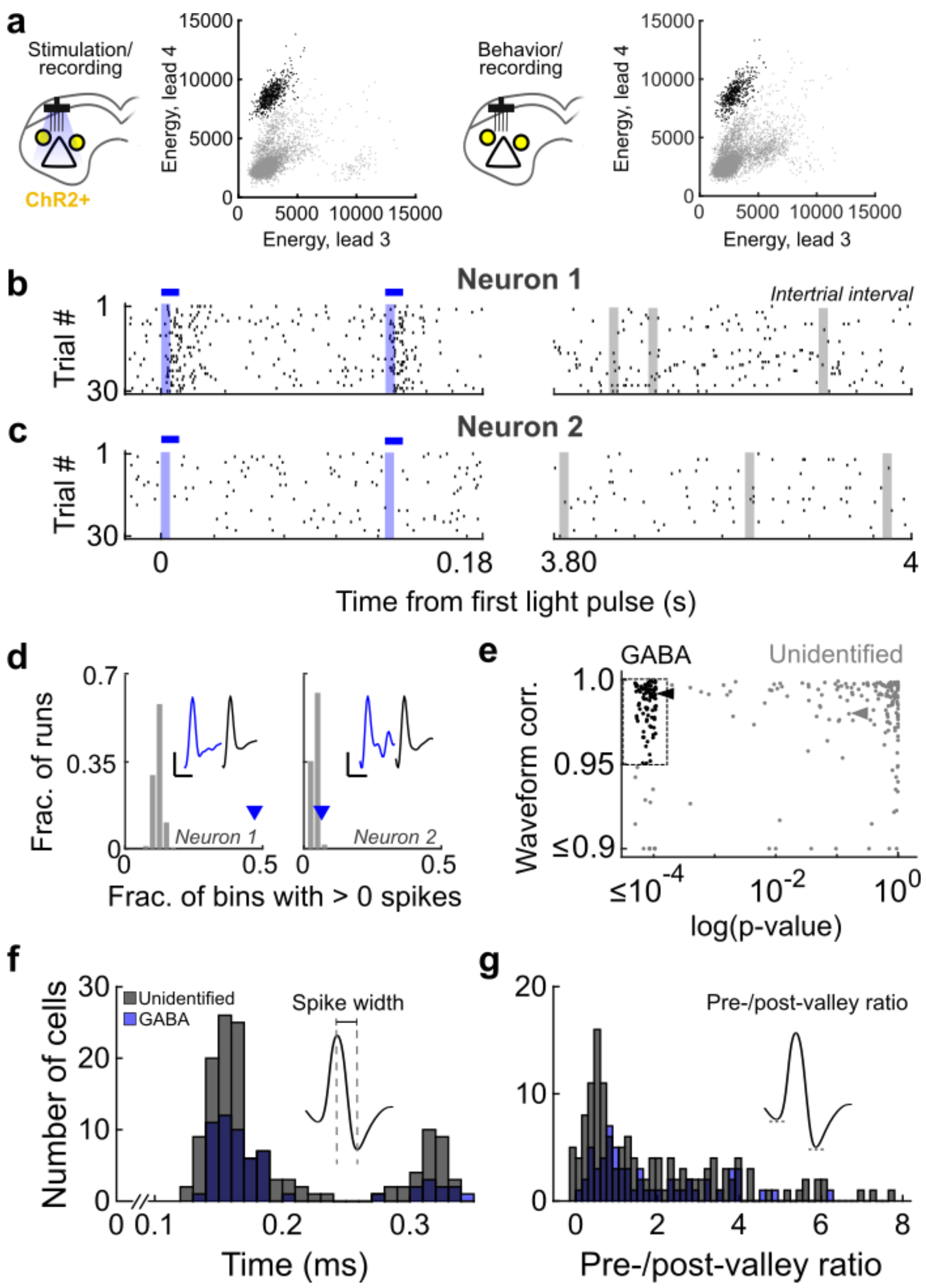


**Supplementary Figure 3: Optogenetic identification of GABAergic SC neurons.**
(**a**) Overview: In an optotagging session performed in conjunction with behavior, we identified neurons with consistent low-latency responses to light (left; see **b-e**). Neurons were “tracked” during the behavioral session (right) based on clustering of waveform features. Energy is the square root of the sum of the squared voltage, reflecting the height and depth of the waveform peak and valley, respectively. Each symbol shows 1 spike waveform; black corresponds to waveforms of identified GABAergic neuron. (**b**) Example rasters for 1 neuron in optotagging session during light-on (left) and light-off periods (inter-trial interval, right). Optotagging sessions consisted of at least 50 trials (only 30 trials shown) of 8Hz light delivery (10 ms light-on/115 ms light-off) followed by 4 s of no light delivery (i.e., inter-trial interval). Analysis of light-induced activity was limited to the first 5 ms

to exclude polysynaptic activity. Blue shading, 5 ms analysis bins during light delivery. Horizontal blue lines show total duration of light pulses (10ms; only 2 of 10 per trial shown). Gray shading, randomly sampled 5 ms analysis bins during the light-off period for 1 run of the simulation (only 3 of 10 per trial shown). (**c**) As in **b**, for a second example neuron. (**d**) Left: Gray: For neuron shown in **b**, frequency (across 5000 simulation runs) of the fraction of light-off analysis bins that contain > 0 spikes. Blue arrowhead, fraction of light-on analysis bins that contain > 0 spikes, which differed from light-off distribution ($p < 0.0002$, permutation test). Insets, mean spike waveforms during light-on periods (i.e. light-driven, blue) and light-off periods (i.e., spontaneous, black) from 1 tetrode lead (waveform correlation = 0.992). Scale bars, 25 µV by 500 µs. Right: As described, for neuron shown in **c**. Waveform correlation = 0.976. (**e**) GABAergic neurons were identified based on a low probability that the measured fraction of light-on analysis bins containing > 0 spikes occurred by chance ($p < 0.0002$, vertical dashed line) and a high correlation between mean spike waveforms during light-on and light-off periods ($r^2 > 0.95$, horizontal dashed line). Each symbol corresponds to 1 neuron (black, identified GABAergic; gray, unidentified). ◄, neuron shown in **b**; ◄, neuron shown in **c**. (**f-g**) Neurons were not identifiable as GABAergic based on waveform features alone; e.g., spike width (**f**) or pre-valley to post-valley ratio (**g**). All other features yielded similar results to **g** and **f**.

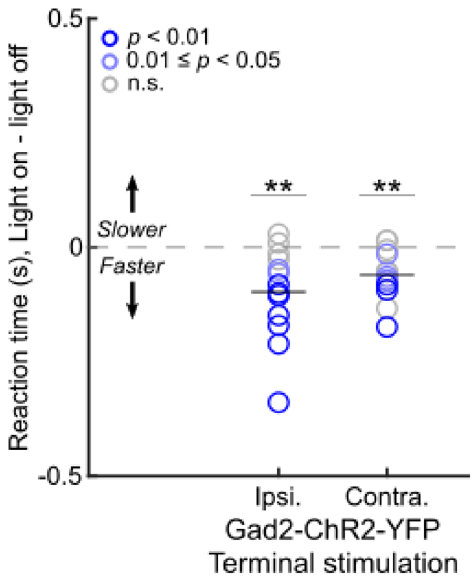


**Supplementary Figure 4: Effect of exciting intercollicular terminals of GABAergic SC neurons on reaction times.**
On each trial, reaction time was calculated as the duration between go signal and odor port exit. In each session, median reaction time was calculated separately for light-on and light-off ipsilateral and contralateral trials, and the difference in medians within each direction is shown. Each session is represented by 2 symbols (1 for ipsilateral trials and 1 for contralateral trials). Horizontal lines show median within group for each direction (with respect to the somata with photoactivated terminals). Photoactivation shortened reaction times for trials in both directions (light on – light off: Ipsilateral, ** $p = 0.0047$, two-tailed one-sample t-test; Contralateral, ** $p = 0.0019$, two-tailed one-sample t-test).

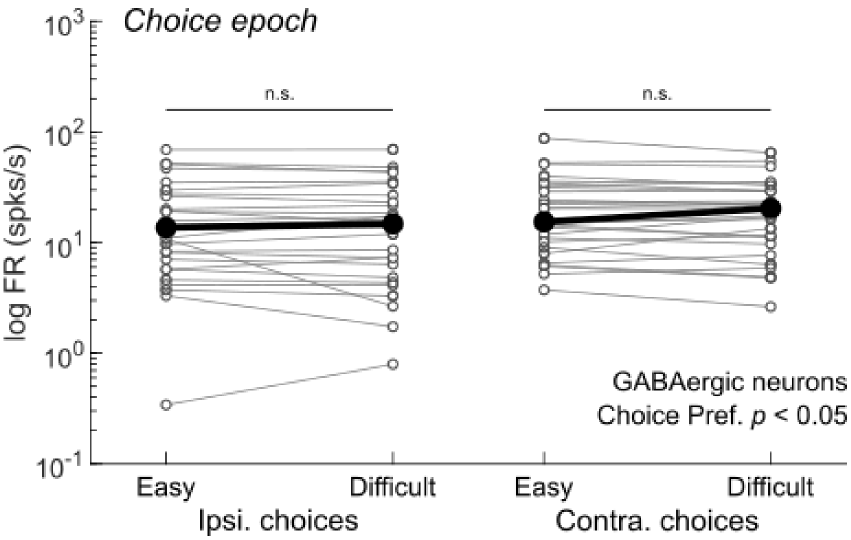


**Supplementary Figure 5: Dependence on trial difficulty of endogenous activity of GABAergic SC neurons during spatial choice.**
Activity of GABAergic neurons with a significant choice preference ($p < 0.05$; black bars in Fig. 4**c**, bottom; n = 29) during the choice epoch on easy (% Ipsi. odor = 5, 20, 80, or 95) and difficult (% Ipsi. odor = 40, 50, or 60) trials shown separately for ipsilateral (Ipsi.) and contralateral (Contra.) choices. Each neuron is represented by a connected pair of small symbols. Large symbols show medians. Activity did not depend on difficulty (Ipsi. choices, $p = 0.642$, two-tailed Mann-Whitney U test; Contra. choices, $p = 0.5963$, two-tailed Mann-Whitney U test).

| Mouse | Stimulation/behavior (# of sessions) | Stimulation/recording (# of neurons) | Behavior/recording (# of neurons) | Stimulation/behavior/recording (# of neurons) | GABAergic terminal stimulation (# of sessions) |
|---|---|---|---|---|---|
| **Abw25** | **12** | **12** | **12** | — | — |
| **Abw35** | **10** | **35** | **29** | — | **3** |
| **Abw36** | **7** | — | — | — | **1** |
| **Abw42** | **8** | **33** | **29** | — | — |
| **Abw43** | **8** | **31** | **16** | — | — |
| Abw49 | 8 | 44 | 42 | 9 | — |
| Abw53 | 10 | 38 | 33 | 12 | 4 |
| Abw57 | 8 | 38 | 36 | 14 | 3 |
| Abw61 | 8 | 29 | 21 | 16 | — |
| Abw63 | 8 | 22 | 8 | 12 | — |
| Abw67 | 9 | 19 | 13 | 2 | 2 |
| Total | 96 sessions | 301 neurons | 239 neurons | 65 neurons | 13 sessions |

**Supplementary Table 1: Mice used in the identification and analyses of GABAergic SC neurons**
